# A co-culture microplate for real-time measurement of microbial interactions

**DOI:** 10.1101/2021.01.07.425753

**Authors:** Charles Jo, David B. Bernstein, Natalie Vaisman, Horacio M. Frydman, Daniel Segrè

## Abstract

The dynamic structures of microbial communities emerge from the complex network of interactions between their constituent microbial organisms. Quantitative measurements of these microbial interactions are important for understanding and engineering microbial community structure. Here, we present the development and application of the BioMe plate, a redesigned microplate device in which pairs of wells are separated by porous membranes. BioMe facilitates the measurement of dynamic microbial interactions and integrates easily with standard laboratory equipment. We first applied BioMe to recapitulate recently characterized, natural symbiotic interactions between bacteria isolated from the *D. melanogaster* gut microbiome. Specifically, the BioMe plate allowed us to observe the benefit provided by two *Lactobacilli* strains to an *Acetobacter* strain. We next explored the use of BioMe to gain quantitative insight into the engineered obligate syntrophic interaction between a pair of *E. coli* amino acid auxotrophs. We integrated experimental observations with a mechanistic computational model to quantify key parameters associated with this syntrophic interaction, including metabolite secretion and diffusion rates. This model also allowed us to explain the slow growth observed for auxotrophs growing in adjacent wells, by demonstrating that under the relevant range of parameters, local exchange between auxotrophs is essential for efficient growth. The BioMe plate provides a scalable and flexible approach for the study of dynamic microbial interactions.

## Introduction

The structures of microbial communities, their composition, diversity, and stability, are emergent properties shaped by the interactions between their constituents (1–5). There are many possible modes of interaction between microorganisms. Direct interactions, which require cell-cell contact, include mechanisms like bacterial conjugation, contact-dependent growth inhibition systems, and intercellular nanotubes (6–9). Microorganisms may also interact via diffusion-mediated mechanisms, such as competition for shared nutrients, production of toxins or communal resources, modulation of environmental conditions, quorum sensing, plasmid exchange, and metabolic cross-feeding (10–14). These multi-faceted interactions within a microbial community have a profound impact on its composition, diversity, and stability (2, 4, 15–17). An appreciation for, and improved understanding of, microbial interactions can elucidate the metabolic and ecological principles of microbial community assembly, as well as facilitate the rational design of novel microbial consortia.

While several approaches have been developed to measure different kinds of inter-microbial interactions, the task of determining the dynamic effect of one microbe’s growth on another remains a challenging task (2, 3, 18–20). Mixed co-cultures, where distinct microbes are cultured in the same vessel, are commonplace. However, measuring the abundance of individual members in co-culture is generally difficult and laborious, and requires sampling for CFU counting (5), or imaging, which is usually restricted to fluorescently labelled strains (21). Plating assays are also prevalent, where a microbe is co-cultured on, or in proximity to, a colony of another microbial species (22–24). Conditioned media assays can also be used to determine how the metabolic byproducts and environmental modifications of one microbe affect the growth of another (4, 25). However, plating assays are limited by their necessity for phenotypic differentiation between microbes and inadequate quantitative readout, whereas conditioned media assays are restricted to the study of unidirectional, non-concurrent interactions. Metagenomic and 16S amplicon sequencing have seen increasing usage to measure relative changes in species abundance, but are ultimately restricted by their cost, temporal resolution, and operational challenges (26, 27).

To address these methodological limitations, systems and devices have been developed to measure co-cultured microbial growth (28). High-throughput microfluidics and micro-droplet based approaches have been designed to screen large numbers of multi-species microbial interactions (29–33). While these approaches offer exciting potential, they can be difficult to apply using standard laboratory equipment, require extensive training and expertise (limiting their accessibility), and often do not allow for the simultaneous observation of the growth of each co-culture member. Alternatively, co-culture systems that utilize a porous membrane to physically isolate individual cultures, and to allow for the exchange of diffusible molecules, have been designed to enable concurrent growth and measurements of distinct microbial cultures (34–37). While these systems restrict microbial interactions to those mediated by diffusible molecules, they have shown promise for culturing previously uncultivated organisms and simultaneously measuring individual growth dynamics in microbial co-cultures. Despite the promise of existing membrane based systems, improvements in their throughput, ease of manufacturing, and ability to seamlessly integrate with common laboratory equipment could enhance their use for the measurement of microbial interactions.

In this work, we present the BioMe plate, a 96-well microplate-based co-culture laboratory device developed to facilitate the observation of dynamic microbial interactions. BioMe separates interacting microbial cultures with a porous membrane, physically isolating individual cultures while allowing for the exchange of diffusible molecules. The growth dynamics of interacting microbial cultures can be measured independently using spectroscopic methods of standard laboratory plate readers. The BioMe plate provides a high-throughput 96-well format and a 3D printed design that facilitates manufacturing and modifications.

We first used BioMe to observe a known symbiotic interaction potentially relevant to the microbiome of a host organism. We chose to study a recently reported symbiosis between organisms of the genera *Acetobacter* and *Lactobacillus* isolated from the gut of a *Drosophila melanogaster* laboratory stock (38). The *D. melanogaster* gut microbiome is relatively simple, hosting low bacterial diversity (1–30 species). *Acetobacter* and *Lactobacillus* are the most commonly found bacterial genera in both lab-reared and wild *D. melanogaster* flies, affecting development, metabolism, and behavior (39–41). The presence of these two genera has been suggested to contribute to deterministic processes related to the assembly of the *D. melanogaster* microbiome (42), as they interact through the exchange of metabolic waste products from *Lactobacillus* to *Acetobacter* (38). Here we use BioMe to implement pairwise co-cultures between *A. oryzifermentans* and two distinct *Lactobacillus* strains (*L. plantarum* and *L. brevis*) isolated from lab-reared *D. melanogaster* stocks (43) providing additional insight into their symbiotic interactions.

We next sought to demonstrate the use of the BioMe plate to facilitate the quantitative investigation of a reduced, well-controlled microbial interaction. Specifically, we used the BioMe plate to study the syntrophic interaction between two *E. coli* amino acid auxotrophs. Engineered auxotrophic bacteria have been used as compelling model systems for the study of microbial interactions due to their well characterized metabolic requirements (44–46). In the work of Mee *et al.* (2014) *E. coli* strains were genetically recombineered for a unique single amino acid auxotrophy and then grown together in mixed co-cultures to identify syntrophic pairs where distinct amino acid auxotrophs could sustain each other’s growth. A syntrophic pair of *E. coli* auxotrophs from this study, the lysine and isoleucine auxotrophs (referred to as ΔLys and ΔIle in the text), were co-cultured in the BioMe plate to further characterize and quantify the syntrophic interaction between these two auxotrophs.

In parallel to the device itself, we developed a computational model of the growth and nutrient dynamics of two strains grown in connected BioMe wells. We applied this model to the syntrophic *E. coli* interaction to determine the factors that underlie their co-dependent cross-feeding relationship. In particular, we used this model, alongside an approximate Bayesian computation based approach, to infer plausible ranges for interaction parameters related to the amino acid’s diffusion across the membrane and leakage out of *E. coli* cells. Together, these efforts demonstrate the scope of novel questions and experiments that are enabled by the BioMe plate, ultimately improving our understanding of the metabolic interactions and ecological relationships that shape microbial community structure.

## Results

### BioMe development

We developed a microplate-based co-culture device, the BioMe plate, which enables the quantitative measurement of microbial interactions (Figure 1). A series of vertical porous membranes physically isolate constituent members of pairwise microbial interactions while allowing for the exchange of diffusible molecules. The physical segregation of interacting microbial cultures enables real-time growth dynamics measurements of each microbial population. A range of micro-scale pore sizes can be selected from, depending on the user’s desired application, molecule size selectivity, and degree of diffusion across the membrane.

**Figure 1:**
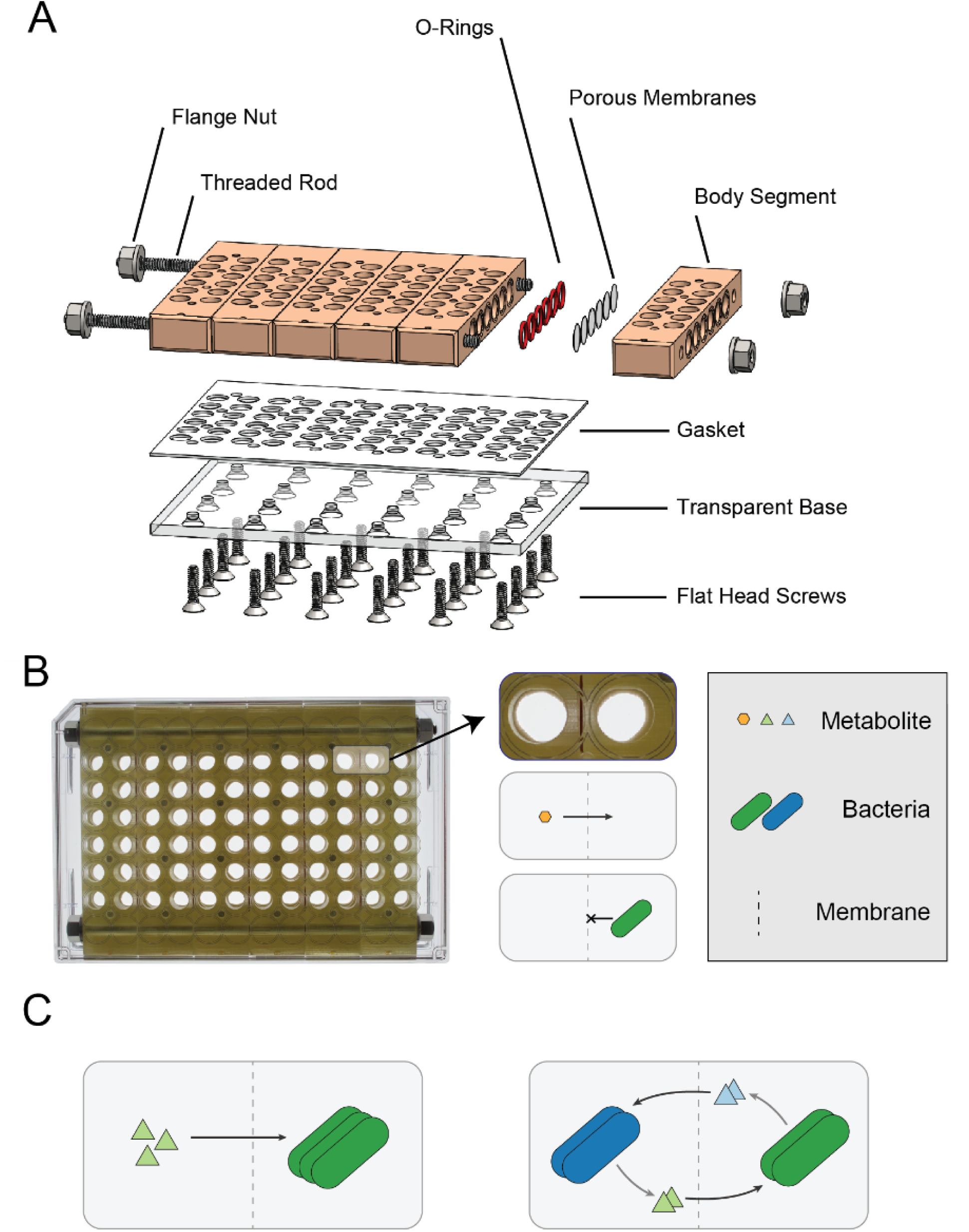
The BioMe Plate. **(A)** Exploded view of the computer-aided design of the BioMe plate. The different components are assembled to form a microplate-based co-culture device. **(B)** Photograph of the fully assembled BioMe plate (top view). Adjacent wells are separated by a porous membrane that allows the diffusion of metabolites and small molecules but not microbes. **(C)** The BioMe plate enables a variety of microbiological experiments. This includes metabolite growth assays (*left*) to observe microbial population growth when seeded across from a metabolite, and co-culture assays (*right*) to characterize and quantify the effects of co-cultured growth between a pair of interacting microbial cultures.

The BioMe plate is comprised of several components, which are further detailed in Supplemental Table 1. The standard 96-well microplate design is fragmented into six discrete body segments, which form the wells of the device. Each well has an opening on the side of the body segment that connects it to its respective co-culture well in a separate body segment, via O-ring and porous polycarbonate membrane. The body segments are laterally fastened and sealed using rods and nuts, yielding a total capacity of 30 pairwise co-culture assays. Of note, the body segments can be machined from polypropylene or stereolithography (SLA) 3D printed using a biocompatible and autoclavable dental resin, circumventing the need for labor-intensive machining. A transparent base forms the bottom of the plate, allowing for real-time spectroscopic readings (optical density or fluorescence) of each well. The base is machined from polycarbonate sheet and is vertically fastened and sealed to the assembled body chassis via a laser-cut gasket and screws. This forms the core of the BioMe plate, which is housed in between two clear 96-well plate lids to prevent evaporation and equipment damage if leakage were to ever occur.

### BioMe is sterilizable, leak-proof, and enables small molecule diffusion

A sterilized BioMe plate can be reused to ensure cost-effective, contamination-free repeated use. The porous membranes are disposable and replaced per use. Leakage tests demonstrated a water-tight seal throughout (see Methods, section: Leakage Test) and sterilization validation tests demonstrated successful de-contamination of the BioMe plate (see Methods, section: Sterilization Protocol and Validation). The ability of various membrane pore-sizes to restrict crossover of microbial cells was tested; crossover was observed in several experiments, especially for larger pore sizes (0.4μm) and rarely for smaller pore sizes (0.03, 0.1, 0.2μm) (see Results, section: Syntrophic *E. coli* Amino Acid Auxotroph Interaction, and Discussion).

Small molecule diffusion across the porous membranes was tested using colorimetric assay dyes. Specifically, the BioMe plate was loaded with two different dyes, phenol red and bromocresol purple, and their diffusion across membranes with varying pore sizes was monitored (Figure 2A and S2.1). Phenol red and bromocresol purple are typically used as pH indicators, but when measured at their isosbestic points, 478nm and 490nm, respectively (47, 48), their concentrations can be inferred from optical density (absorbance measurements). We developed calibration curves for each dye to ensure accurate extrapolation of concentration from optical density (Supplemental Figure S2.2). Both dyes were observed to diffuse across the porous membrane with a diffusion rate that increased monotonically for increasing pore size, and were not observed to diffuse across a membrane with no pores. The raw optical density data for the diffusion and calibration experiments, laid out as they were on the BioMe plate, are shown in supplemental Figures S2.3 and S2.4.

**Figure 2:**
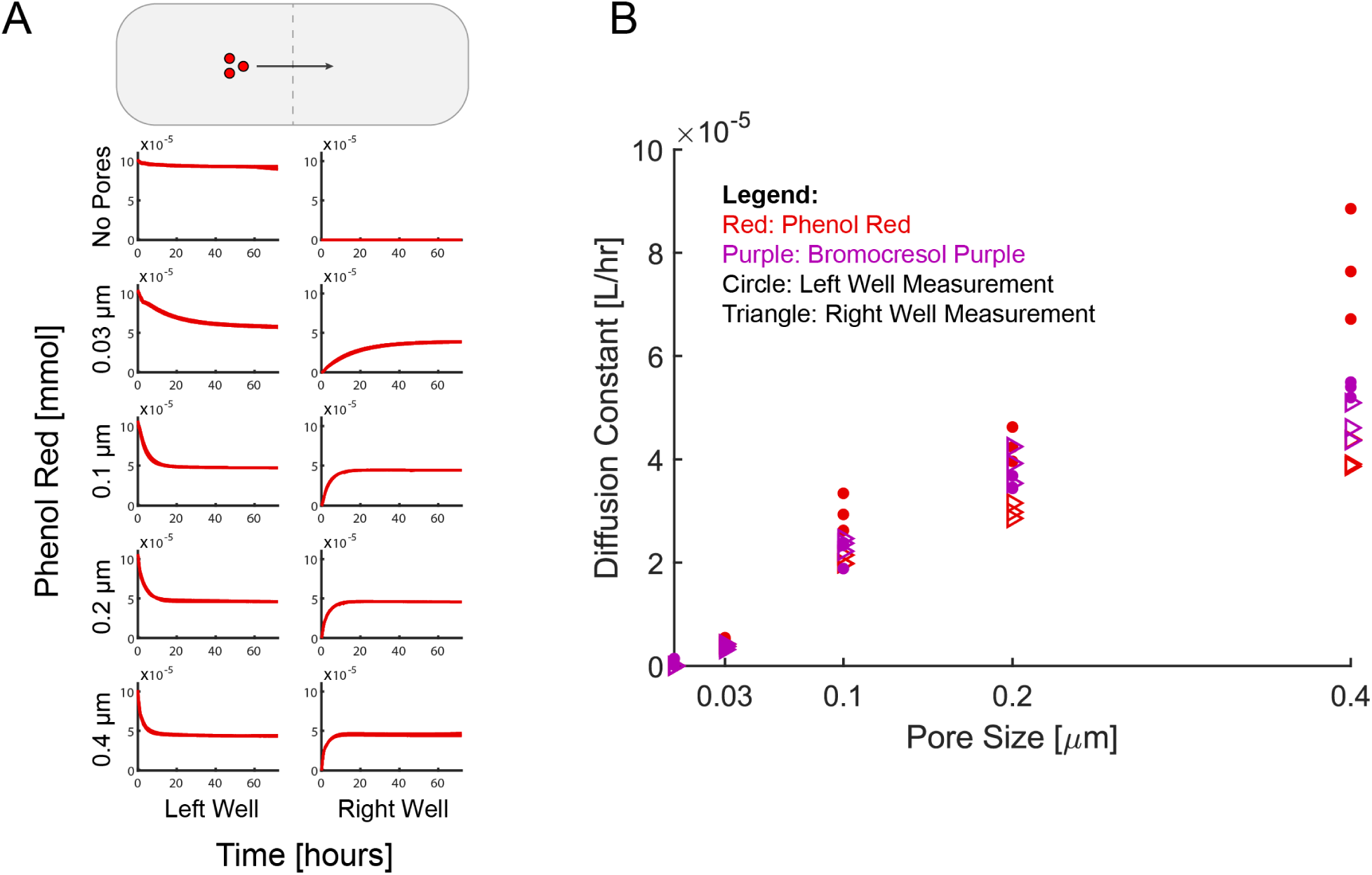
Chemical Dye Diffusion Across Membranes with Variable Pore Size in the BioMe Plate. A small-molecule dye (400μM concentration of phenol red or bromocresol purple) was loaded into columns of wells to the left of the membrane in the BioMe plate, for varying membrane pore sizes: no pores, 0.03, 0.1, 0.2, and 0.4μm. Optical density at isosbestic point (478nm for phenol red, 490nm for bromocresol purple) was measured for 72hrs, in 15min intervals. Dye concentration was inferred by linear calibration to optical density. **(A)** Time course of calibrated dye concentration for phenol red with membranes of different pore sizes. Replicates of n=3 (all overlapping curves are shown on the plot) were conducted for each pore size. **(B)** Diffusion constants for phenol red and bromocresol purple for varying membrane pore sizes, calculated by fitting to dye diffusion model using data from both the left and right well measurements.

Diffusion coefficients of all tested membrane pore sizes for each chemical dye were inferred using a gradient-driven diffusion model (further described in the *Computational Modeling* section). This model gives rise to an exponential function describing the time-dependent concentration of the molecules in either well of the system, as the dyes diffuse across the membrane (Supplemental Equations S1). An exponential function was fit to the dye diffusion data (See Methods and Figure S2.5) to infer a range of estimates for our model’s diffusion constant for both dyes and various pore sizes: 0.03, 0.1, 0.2, and 0.4μm. As shown in Figure 2B, the mean diffusion constant increases monotonically and non-linearly with increasing pore size. Interestingly, the estimated diffusion constant was consistently higher when calculated using the curves from the well where the dye was initially placed as opposed to when using the curves from the opposite well into which the dye diffused and this effect was more pronounced for the larger pore sizes and for phenol red (see Discussion for additional details).

### Symbiotic interaction between *Drosophila melanogaster* gut microbiome species

As a first biological application of the BioMe plate, we confirmed a recently characterized interaction among specific genera of bacteria from the *Drosophila melanogaster* gut microbiome. In particular, we focused on three distinct bacterial strains previously isolated from laboratory-bred flies: one species of *Acetobacter* (*A. oryzifermentans*) and two species of *Lactobacillus* (*L. plantarum and L. brevis*) (43). The BioMe plate was used to test and observe every pairwise co-culture combination between these three strains. Additionally, relevant controls were included in the experiments, i.e. organisms paired with themselves (self-control) and organisms paired with wells containing growth medium but no other organism (media-control) (Figure 3, Supplemental Figure S3.1 and S3.2). A 0.03μm pore size membrane was chosen to allow exchange of metabolites but ensure no microbial cross-over, as done for the iChip device (49).

**Figure 3:**
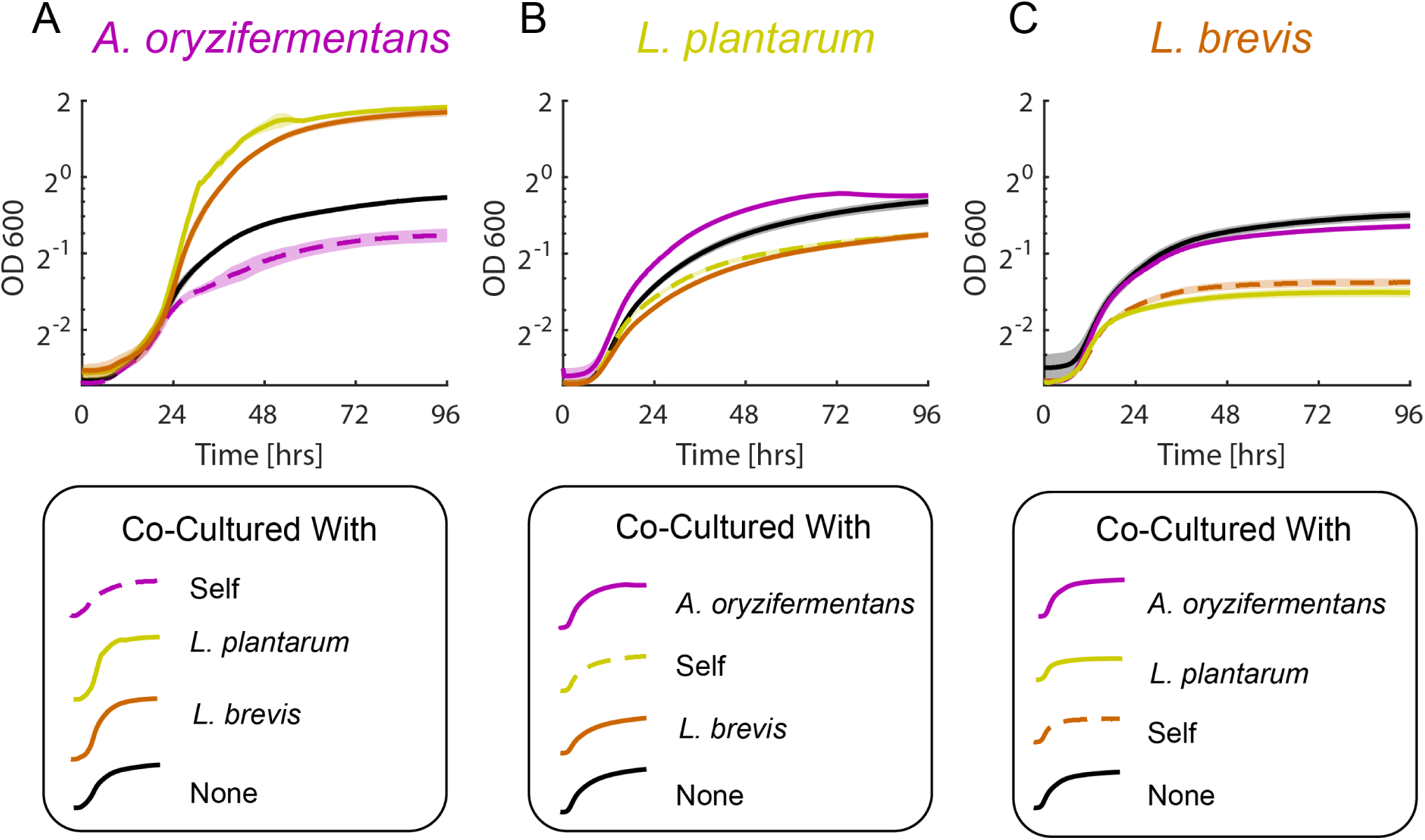
Commensal Relationship between *Drosophila melanogaster* Gut-Associated Microbes. Pairwise co-cultures between all combinations of *Acetobacter oryzifermentans* **(A)**, *Lactobacillus plantarum* **(B)**, and *Lactobacillus brevis* **(C)**. Growth curves are plotted on a log base 2 scale. Membrane pore size of 0.03μm was used throughout these experiments. Lines indicate mean growth curves and shaded regions indicate standard error (n=4 for cross-species co-cultures, n=3 for media-control “none”, n=6 for self-control “self” for *L. plantarum* and *L. brevis* and n = 4 for “self” for *A. oryzifermentans* cultures as two outliers were left out, see supplemental figures S3.1 and S3.2 for additional details).

The co-culture experiments revealed an increased growth of *A. oryzifermentans* when co-cultured with either *Lactobacillus* strain, demonstrating a clear benefit relative to its growth when co-cultured with itself or no microbe in the opposing well. Interestingly, for both *L. brevis* and *L. plantarum*, their growth in co-culture with the *Acetobacter* strain was similar to their growth in respective media-controls, but still demonstrated improved growth relative to both their self-controls and co-culture with the other *Lactobacillus* strain. Our results, uniquely enabled by the BioMe plate, suggest that the *Acetobacter* population disproportionately benefits from this metabolite-mediated interaction. BioMe could serve as the starting point for further exploration of this cross-species interaction, complementing existing approaches (38, 50, 51).

### Syntrophic interaction between *E. coli* Amino Acid Auxotrophs

Next, we used BioMe to enable a more in-depth and quantitative study of a model microbial interaction in which two strains have been engineered to exchange essential metabolites in order to grow. Specifically, we studied the syntrophic interaction between a pair of *E. coli* amino acid auxotrophs: ΔLys and ΔIle. These strains were engineered to be unable to biosynthesize lysine or isoleucine, respectively, requiring supplementation of the missing amino acid to grow in monoculture. Despite their inability to grow in monoculture without supplementation, both strains are able to grow when placed together in a mixed co-culture (44).

The first test of this system in the BioMe device was a validation that amino acids could diffuse across membranes of various pore sizes and support the growth of an auxotrophic *E. coli* strain in the adjacent well. Figure 4 shows an example of this validation for lysine diffusion with the 0.1μm pore size. Lysine was shown to diffuse across the membrane, with comparable growth to the positive control where lysine was added into the same well as the microbial culture (Figure 4A and B). A negative control experiment, lacking the supplemental amino acid, displayed no growth, and thus confirmed the amino acid auxotrophy (Figure 4C). A second negative control experiment, displaying lack of growth for membranes with no pores, confirmed the integrity of the device seal (Figure 4D). These results were repeated with ΔIle and isoleucine, and for varying pore sizes: no pores, 0.03, 0.1, 0.2, and 0.4μm. We found similar results in all cases, with comparable growth between diffusion and positive control conditions and no growth in either of the negative controls (Figure S4.1, S4.2, and S4.3).

**Figure 4:**
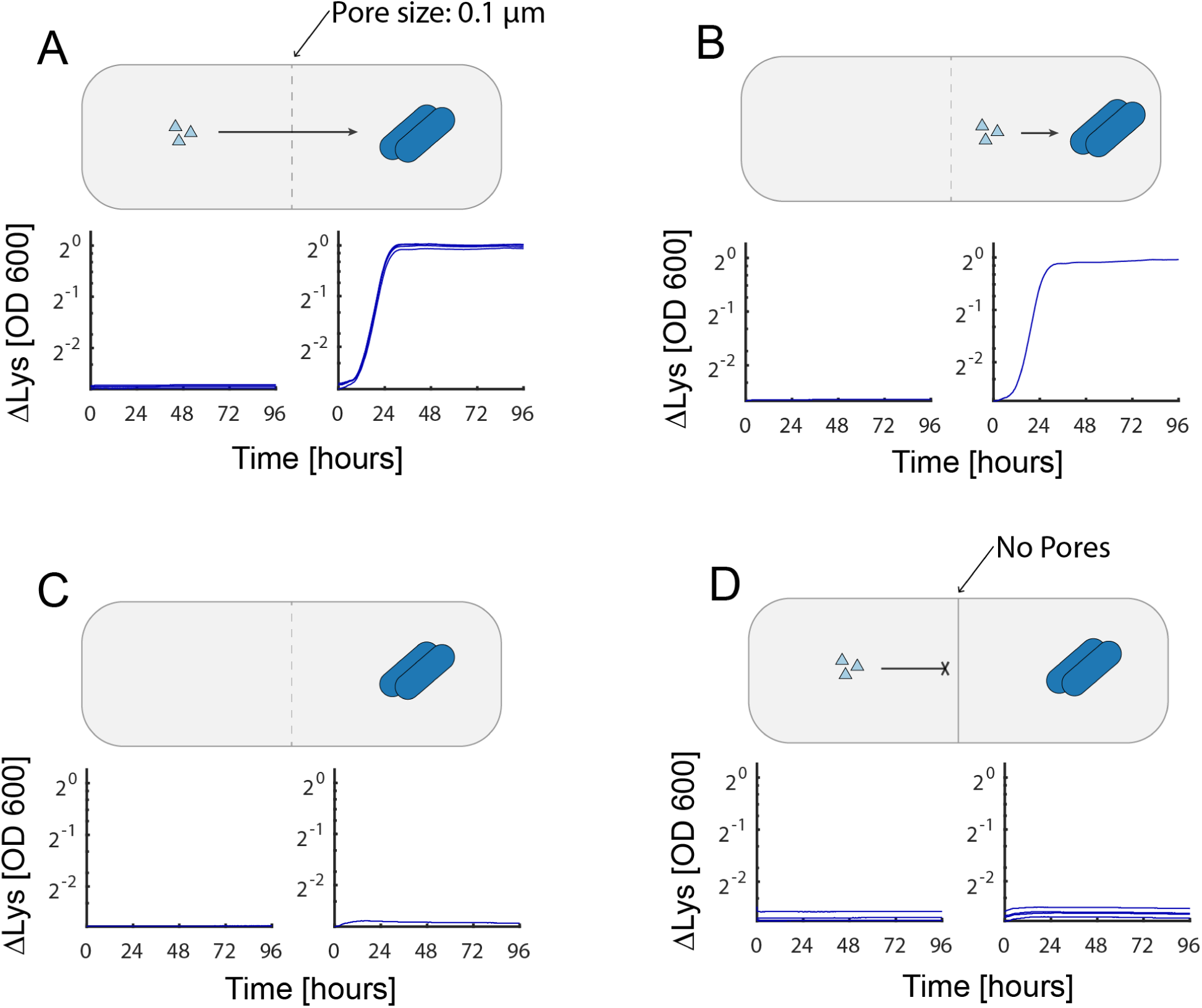
Lysine Diffusion Bioassay with *E. coli ΔLys* in the BioMe Plate. Various amino acid diffusion conditions were tested in the BioMe plate. **(A)** ΔLys *E. coli* auxotrophs grown in minimal media with supplemental lysine across the 0.1μm membrane (n=4). **(B)** Positive control with lysine supplemented in the same well as the microbial culture (n=1). **(C)** Negative control with no supplemental lysine provided (n=1). **(D)** Negative control with supplemental lysine provided across a membrane with no pores (n=4).

We next co-cultured ΔLys and ΔIle in the BioMe plate to measure the dynamics of their metabolic interaction (Figure 5). As expected, each strain can complement, to some extent, the amino acid missing in the other, confirming the existence of cross-feeding. Surprisingly, however, the auxotrophs grew significantly slower when inoculated in paired wells separated by a porous membrane than when grown in the same well (Figure 5A and B). The membrane significantly impeded the growth of both members in the syntrophic interaction. As shown in Fig. 5A, this effect is asymmetric, in that ΔIle seems to be able to help ΔLys more than ΔLys helps ΔIle, though the growth of ΔIle is still higher than the negative control. Negative controls further confirmed that neither auxotrophs could grow without the partner strain (Figure 5C and D) or when separated by a membrane with no pores (Figure 5E). Qualitatively similar results were observed for membrane pore sizes of 0.03, 0.1, and 0.2μm (Figure S5.1 and S5.2). For membranes of 0.4μm pore size, microbial cross-over through and/or around the membrane/O-ring junction occurred in all replicates (measured by selective plating at the end of the experiment, Supplemental Table 3), preventing any measurement of interaction dynamics. The slow-down of cross-feeding across the membrane (Fig. 5) seems curious, especially in light of the fact that amino acid diffusivity seemed not to be an issue when each organism had the auxotrophic amino acid on the other side of the membrane (Fig. 4). This observation pointed to a subtler concentration-dependent effect, and we wondered whether the quantitative nature of the BioMe experiments could help us gain a deeper understanding of the dynamics underpinning this phenomenon.

**Figure 5:**
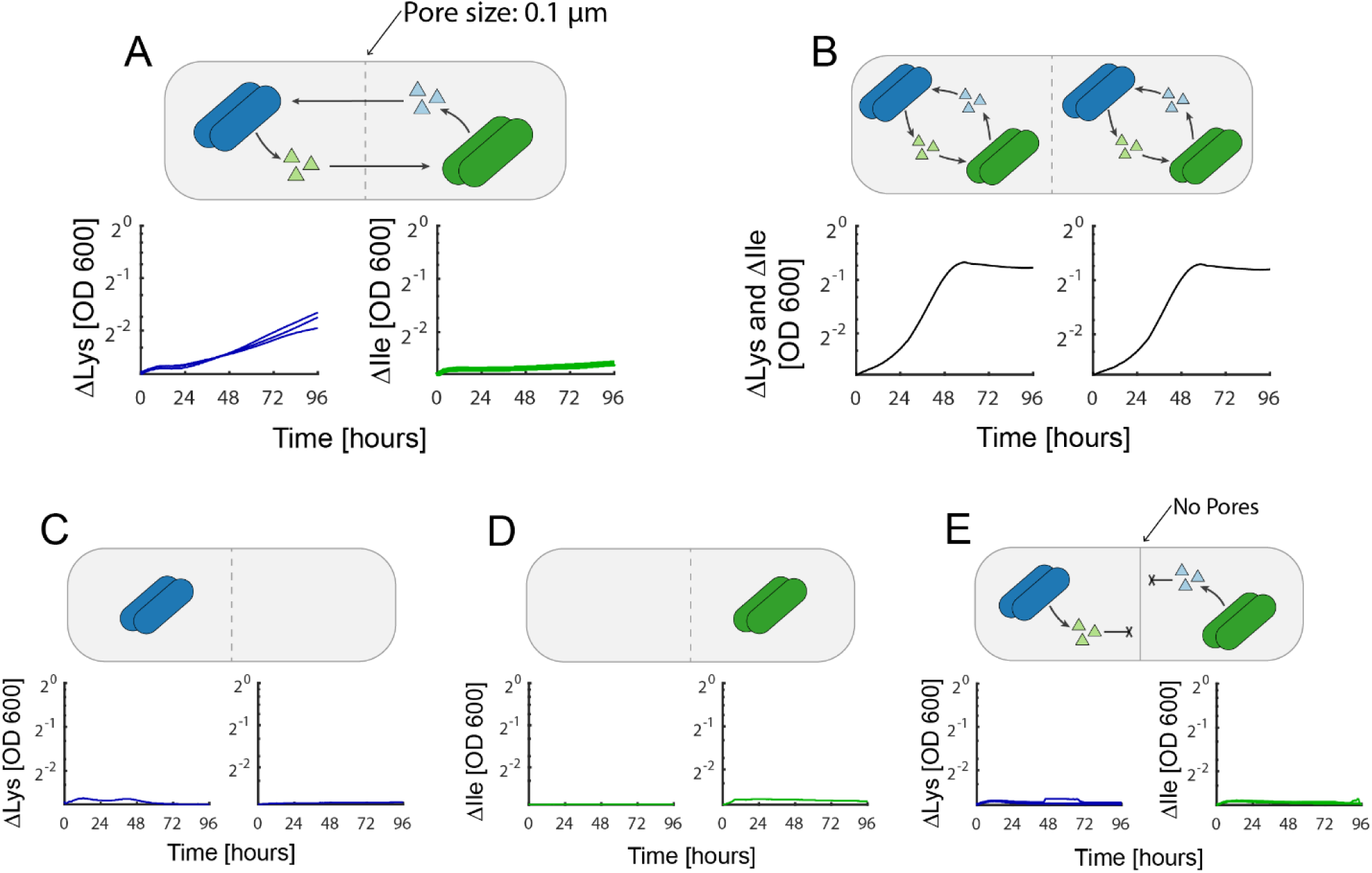
Syntrophic *E. coli* Lysine and Isoleucine Auxotroph Co-Culture. Mutualistic cross-feeding between ΔLys and ΔIle amino acid auxotrophs. **(A)** ΔLys (blue) and ΔIle (green) grown across 0.1µm porous membrane (n=3). **(B)** Positive control with ΔLys and ΔIle grown in the same wells (n=1). **(C&D)** Negative controls with ΔLys and ΔIle without a syntrophic partner, respectively (n=1). **(E)** Negative control with ΔLys and ΔIle separated by a membrane with no pores (n=3).

### Computational modeling of BioMe co-cultures provides insight into interaction parameters

To further investigate the interaction dynamics observed experimentally with the BioMe plate, we developed a computational model describing the syntrophic *E. coli* auxotroph co-culture experiments. This model captures the dynamics within two interacting wells of the BioMe plate, by simulating the processes of metabolite diffusion across the membrane, glucose and amino acid uptake by the *E. coli* auxotrophs, and a stoichiometric leakage of the respective partner’s auxotrophy amino acids (Figure 6); the full list of equations and parameters is available in supplemental Equations S2 and at https://github.com/segrelab/co_culture_device. Using this computational model, all performed experiments involving the *E. coli* amino acid auxotrophs can be simulated by changing the initial conditions of the model to match those of the experiment (Figure S6.1).

**Figure 6:**
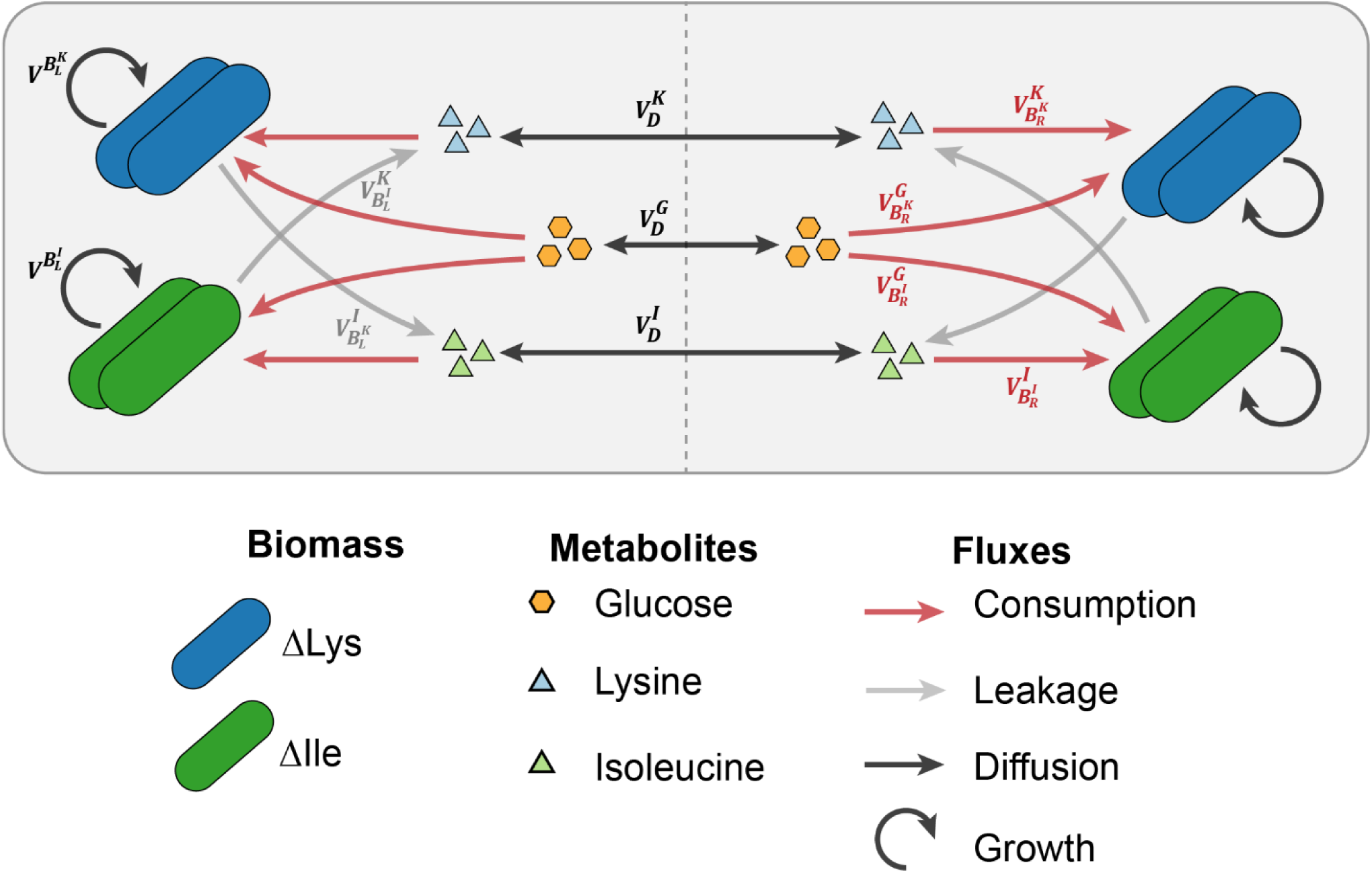
Computational Model of Syntrophic *E. coli* Auxotroph Growth Dynamics in BioMe. The fluxes of metabolites simulated by the model are illustrated in this figure, including: the diffusion of glucose and amino acids across the porous membrane (black arrows), the uptake of glucose and auxotrophy amino acids supporting growth (red arrows), and the leakage of the respective partner’s auxotrophy amino acids (gray arrows). The growth of each auxotroph’s biomass is also represented (circular arrow). Supplemental Equations S2 describe all parameters and the full dynamics of the system.

We integrated this computational model with our syntrophic co-culture experiment to explore the parameter space that gives rise to our primary qualitative observation – that both members of the mutualistically cross-feeding pair grew significantly faster when in the same well than when interacting across the membrane. The parameters for uptake kinetics and biomass stoichiometry were estimated from literature (Supplemental Equations 3). The parameter space was then explored for two important parameters of the interaction, which were difficult to estimate from the literature: the diffusion rate of the metabolites across the membrane and the leakage stoichiometry of the amino acids. In particular, upon varying these parameters, we compared the 48-hours predicted growth yield of an auxotroph grown in the same well with its partner relative to the yield in membrane-separated co-culture (Figure 7A). Distinct regions were clearly visible in this parameter space, which gave rise to different qualitative simulated results (Figure 7 A and C). One can see that if cellular leakage of the amino acid is too small (Figure 7A, region 3) the same-well and separate-well simulations behave similarly, i.e. neither supports any growth. If both leakage and diffusion are high enough (Figure 7A, region 1), again the two in silico experiments behave similarly, but this time they both lead to increased yield. Notably there is an intermediate region of the parameter space (Figure 7A, region 2) in which leakage is not a limiting step to guarantee cross-feeding, but diffusion between separate wells is. In this region, the two auxotrophs can secrete enough amino acids to support each other, but diffusion through the BioMe membrane slows down the growth process enough to lead to a detectable difference.

**Figure 7:**
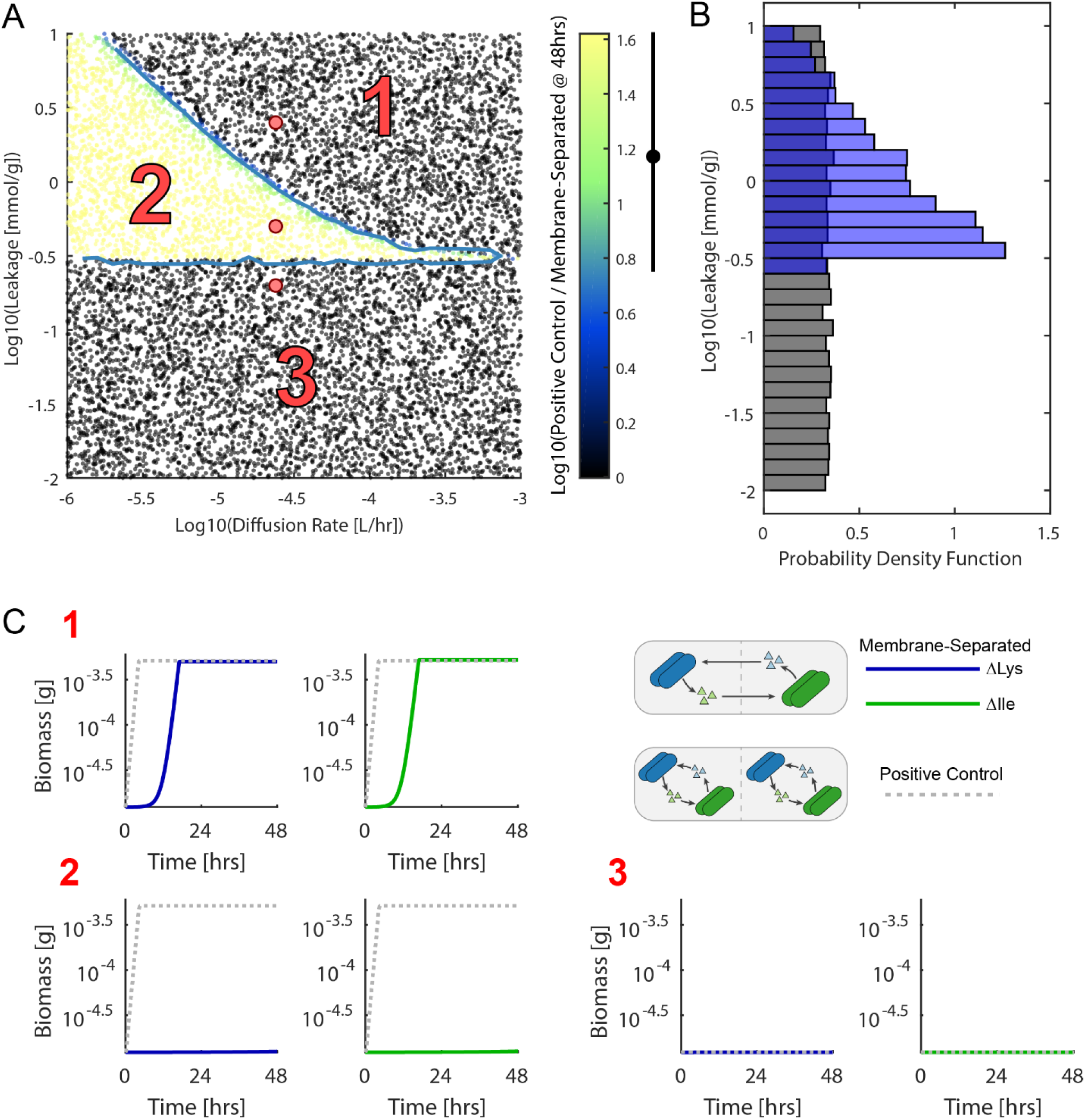
Computational Modeling of *E. coli* auxotroph Co-Culture Interaction Parameters. **(A)** The sampled space of diffusion and leakage parameters is shown. The diffusion rate and leakage stoichiometry were sampled from log uniform prior distributions. Each point represents a different simulation of the model. The color indicates the log base 10 of the ratio between the yield at 48 hours of *E. coli* auxotrophs grown in the same well (positive control) divided by the yield in opposite wells (membrane-separated). This value was used as a statistic for approximate Bayesian computation. The experimental average of this statistic (1.18) and two times the standard deviation (0.448) are indicated by the black dot and bar next to the color bar. Points within this range are separated by a blue line outlining region 2 in the scatter plot. **(B)** The prior (gray) and posterior (blue) distributions of the leakage stoichiometry parameter are plotted as histograms. The prior distribution was sampled from a log uniform distribution. The posterior distribution was calculated by accepting points with a statistic value within 2 standard deviations of the average experimentally observed value. **(C)** Example growth curves for different regions of the parameter space in A are shown. Simulated growth curves of ΔLys (blue line) and ΔIle (green line) grown across from each other (membrane-separated), and grown together (gray dotted line, positive control), are shown for three different regions of parameter space.

The identification of the intermediate region (Fig. 7A, region 2) prompted us explore the possibility of using this modeling framework to provide more insight into the experimentally unknown key parameters. In particular, we used an approximate Bayesian computation approach (see Methods) to infer, and model the uncertainty of, the leakage parameter values that were consistent with the observed differential growth yields in the same-well positive control versus the membrane-separated co-culture (Figure 7 B). The posterior distribution of the leakage parameter, conditioned on this experimental observation, was estimated by using a statistic of the logarithm of the ratio of the biomass yields at 48 hours (Supplemental Equations S4) and an acceptance cut-off of 2 standard deviations around the experimentally measured mean value (See Methods for further details). To further investigate the asymmetric nature of our observations, a similar analysis was conducted where the lysine and isoleucine leakage parameters were sampled from independent prior distributions, and could thus be inferred separately (Supplemental Figure S7.1). This gave rise to a lysine leakage parameter that was higher than that of isoleucine, consistent with our observation of increased ΔLys growth. Furthermore, noise was added to the prior distributions of the literature inferred parameters, by sampling them from a log uniform distribution spanning one order of magnitude around their literature inferred values (Supplemental Figure S7.2 & S7.3). Even in the presence of this amount of noise in these values, the leakage stoichiometry parameter was effectively constrained by the data. Through this analysis, we integrated novel observations obtained with the BioMe plate with a computational model to infer plausible ranges of the leakage parameter governing the interaction between these auxotrophic *E. coli* mutants. The posterior distributions obtained from this approximate Bayesian computation approach represent these inferences and their uncertainties, given the assumptions of our model and the data that we conditioned it on. A similar approach could be applied to any observations of interaction dynamics from the BioMe plate to gain insight into how these observations further constrain the inference of different parameters of interest.

## Discussion

We utilized the newly developed BioMe plate to observe a natural symbiotic interaction between organisms found in the *D. melanogaster* gut microbiome, and to quantitatively study the interaction between two syntrophic engineered *E. coli* mutants. While our *D. melanogaster* results demonstrate the utility of BioMe to study a natural microbial interaction between two non-model organisms, corroborating recently published results (38), the better characterized *E. coli* auxotroph interaction allowed us to integrate the data obtained with BioMe into a computational model providing quantitative insight into the parameters of this interaction.

Moving forward, as computational modeling of microbial physiology advances, it may be possible to apply such modeling frameworks more ubiquitously. For example, genome-scale metabolic models could be incorporated into a spatial model of the BioMe device to represent the metabolic processes occurring in each interacting population and model emergent interactions (11). Such a model could be used, alongside BioMe observations, to gain quantitative, genome-scale insight into the interactions between microbial organisms, where the mechanism of interaction is not known *a priori*.

Our investigation of the *E. coli* lysine and isoleucine auxotrophs revealed that their syntrophic interaction was altered by the presence of a porous membrane separating the interacting auxotrophs. These results suggest that certain “high-stakes” interactions could be localized, such that highly-proximal ecological neighbors reap the majority of the benefits from “leaked” communal resources. Leveraging a mechanistic, dynamical model for the syntrophic *E. coli* auxotroph co-culture experiment, we inferred plausible leakage parameters that govern this localized interaction. This result corroborates and complements the recent imaging-based finding that pairs of *E. coli* proline and tryptophan auxotrophs can best help each other within a small local radius around each cell (52). While these local interactions may be dominated by diffusion-mediated processes, we cannot rule out, based on our experiment, that cross-feeding is also facilitated by contact-based mechanisms. Similar phenomena were reported, for example, in bacterial-fungal interactions, which were shown to require physical association for the activation of a cryptic biosynthetic pathway of secondary metabolism in the fungus (53). The local nature of these interactions may have important evolutionary and ecological consequences (54, 55), and the BioMe device is a promising platform for the continued study of such phenomena. It is conceivable that analyses similar to the one preformed here for the *E. coli* auxotrophs, could be extended in the future to the study of natural interaction partners, such as the *D. melanogaster* gut microbiome strains (38, 50, 51), or other pairs of strains suspected to interact with each other through the exchange of metabolites.

There are some key limitations to the use of the BioMe plate. First, as is clear from our results, the porous membranes can have a significant impact on mutualistic cross-feeding interactions. In addition to altering the dynamics of metabolite exchange, the membrane limits interactions to only those mediated by diffusible molecules. Despite placing a constraint on the type of interactions that can be investigated, this limitation can be beneficial for the quantitative study and design of microbial consortia. For example, limiting interactions to those mediated by diffusible molecules makes it easier to robustly integrate and compare experimental measurements with metabolic models. Another limitation of the BioMe device is the fact that microbial cross-over can occur across the membranes. In our syntrophic auxotrophs co-culture experiment we observed microbial cross-over in 2/9 pairwise assays with 0.03, 0.1, and 0.2μm pore sizes, and 3/3 for the 0.4μm pore size. These studies were exquisitely sensitive for detecting microbial cross-over as the mutualistic interaction between the two strains would effectively amplify any crossed-over organisms. Future studies could focus on more thoroughly characterizing the cross-over process for a variety of different organisms and conditions. For studies aimed at observing microbial interactions, the choice of membrane pore size should be carefully considered to restrict microbial cross-over and simultaneously allow for desired diffusion of metabolites.

Regarding the manufacturing of the BioMe device, it should be noted that while the body segments can be fabricated using either milling or 3D printing methods, there is a significant trade-off between precision and flexibility in these two methods. We found that devices fabricated using milling had finer tolerances and were thus less prone to leakage than those that were 3D printed. However, the 3D printing approach facilitates rapid and easy prototyping of alternative plate layouts. Ultimately, users should choose the fabrication technique that best suits their goals.

Regarding the pore size dependence of small-molecule diffusion in the device, there was an interesting observation in the chemical dye diffusion experiments – the estimated diffusion constant was consistently higher when calculated using data from the input well where the dye was initially placed as opposed to that of the opposite well. This effect was more pronounced for the larger pore sizes and for phenol red. A possible explanation for this effect is that the dye is settling or being sequestered in the membrane/device, and thus appears to be diffusing faster from the initial well and more slowly into the opposite well. This effect was not included in subsequent modeling efforts, as it produced only minor differences in the inferred diffusion constant relative to the range of uncertainty in the diffusion parameter considered in our model.

Although we demonstrate here the use of the BioMe plate to measure pairwise interactions, the device is a flexible technology that could be re-designed to incorporate multi-member assays and higher-order interactions. 3D printing accessibility enables rapid fabrication and validation of potential plate layouts. Going beyond pairwise interactions, selected consortia of microbial organisms could be grown in each well of the BioMe plate to facilitate the investigation of higher-order interactions, which are thought to have an important impact on microbial community structure (56–58). Entire microbial communities could similarly be grown across from individual isolates to observe their growth supporting capabilities, as done with the iChip device (49), but with the added capability of observing growth dynamics. Furthermore, the 3D-printed design of the BioMe body segments could be modified to design novel interaction chambers, such as multiple wells connected to a central mixing chamber or sequential wells connected by porous membranes. Overall, the BioMe plate is relatively easy to manufacture and implement, and the use of this device to study the interaction of microbial organisms from a multitude of different contexts will help improve our understanding of and ability to engineer microbial communities.

## Methods

### BioMe Fabrication

All materials required for BioMe fabrication are detailed in Supplemental Table 1. O-rings, 6-32 flange nuts. 4-40 screws, and membranes were bought ready to use. 6” 6-32 rods were cut to 125mm with a grind wheel. Stock food-grade silicone rubber sheets were laser-cut to gasket specifications using an Epilog Laser Mini 60W laser cutter; wrapping in dampened shop towels helped mitigate charring. The transparent base was CNC milled from clear polycarbonate sheets.

The body segments could either be machined or 3D-printed. Two separate CNC milling operations were required to machine the vertical and horizontal features of the body segment from polypropylene sheet. A drill press was used to complete the spot holes, with each screw hole manually tapped with a 4-40 tap bit, and each body segment deburred and washed.

Alternatively, stereolithography (SLA) resin 3D-printing can be used to fabricate the body segments. FormLabs’ Form2 SLA printers were used with the Dental SG Resin, a biocompatible, autoclavable resin. Screw holes were manually tapped and body segments were sanded to size.

### Leakage Test

A simple visualization test was used to test water-tight seal throughout the assembled BioMe plate. All wells were loaded with 250μL of 100μM phenol red. The core BioMe device was then placed atop a paper towel and fit into the bottom tray and covered with the top lid. The BioMe plate was left in the shaking incubator overnight and no leakage was verified.

### Sterilization Protocol and Validation

To sterilize the BioMe plate after use, the device is disassembled, membranes disposed, components dishwashed, and then autoclaved (Gravity, 30mins exposure/15mins dry). For sterile assembly, a biosafety cabinet is recommended. Membranes are bathed in 70% ethanol for 30mins. Pre-sterilized components are then sequentially assembled with the membranes in sterile conditions. For additional caution, pre-sterilized device components may be ethanol bathed prior to assembly.

Sterilization protocol was validated to ensure no post-contamination. A sterilized re-assembled BioMe plate was loaded with 250μL LB Miller media (10g/L tryptone, 10g/L NaCl, 5g/L yeast extract) per well and placed into a 30°C static incubator for 72hrs. A 10μL aliquot from each well was then plated onto LB agar plates. Plates were then incubated at 30°C for 72hrs. No microbial growth was confirmed.

### Colorimetric Dye Diffusion

Calibration curves for phenol red (PR) and bromocresol purple (BP) were determined to relate optical density at isosbestic point (478nm for PR and 490nm for BP) to concentration. A BioMe plate was assembled with membranes with no pores. Columns were loaded with 250μL of varying concentrations of dye: 450, 400, 350, 300, 250, 200, 150, 100, 50, 0μM; top three rows were used for PR and bottom three rows for BP. Linear fit to OD_isosbestic_ vs. concentration data points used for calibration curve.

Diffusion experiments were used to estimate the diffusion coefficients of various membrane pore sizes for PR and BP. A BioMe plate was assembled with varying membrane pore sizes at each body junction: no pores, 0.03, 0.1, 0.2, and 0.4μm. 250μL of 400μM dye was loaded into the left-column, and 250μL of distilled water in the right-column for each pore size. OD_isosbestic_ was measured for 72hrs, in 15min intervals.

Diffusion constants were estimated using a gradient-driven diffusion model, provided in supplemental Equations S1. The time dependent value on the left hand side of supplemental Equations S1 was calculated from the data and fit to an exponential function (using MATLAB function fit with fittype: exp1) to infer our model’s diffusion constant for both dyes and pore sizes of 0.03, 0.1, 0.2, and 0.4μm. The units for the diffusion constant, *d*, in our equations are L/hr such that the flux has units of mmol/hr. Our model is similar to Fick’s first law, in that there is a gradient driven diffusion, however in our system the diffusion area and distance are fixed and thus lumped into the diffusion parameter giving rise to units that are different from a traditional “diffusion constant”.

### Drosophila Gut Microbiome Interaction

Strains of *Acetobacter oryzifermentans, Lactobacillus brevis,* and *Lactobacillus plantarum* were isolated as previously described (59) and streaked onto YPD agar plates (10g/L peptone, 10g/L yeast extract, 8g/L dextrose, 15g/L agar). Identity and lack of cross-contamination were confirmed by colony PCR using species-specific primers and gel electrophoresis (43). For each strain, four clonal replicates were picked from colonies, and grown in 5mL of YPD broth (10g/L peptone, 10g/L yeast extract, 8g/L dextrose) at 30°C in a static incubator for 20hrs. Cells were centrifuged and pellets washed thrice in PBS. OD600 of 250μL of each culture and PBS blanks were read. Cultures were diluted with YPD media to an OD600=0.1 using following formula:

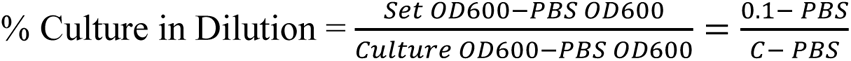

Dilutions were re-done for those not within 10% deviation. Cultures were further diluted 1:100 in YPD media. Assembled BioMe plate, with 0.03μm pores size membranes, was loaded with diluted cultures in appropriate wells and sealed with parafilm. The plate was then run on a plate-reader for 96hrs, 30°C, with OD600 measurements at 15mins intervals.

### Syntrophic Co-Culture Interaction

Diffusion of amino acids through membranes of various pore sizes was validated using Lambda Red-recombineered EcNR1 *E. coli* for knock-out of lysA (ΔLys) and ilvA (ΔIle) (44). Strains were streaked and selected on LB+Cam (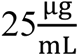 chloramphenicol) agar plates. Strains were grown in 5mL of LB+Cam broth (n=4 for both ΔLys and ΔIle) in a 12-well plate for 24hr inside the plate-reader, with OD600 measurements every 15mins. These growth curves were used to determine dilution time (t_dil_=6hrs) for proceeding experiments (Supplemental Figure S4.4).

For amino acid diffusion experiment, a BioMe plate was assembled with varying membrane pore sizes at each body junction: no pores, 0.03, 0.1, 0.2, and 0.4μm. For either strain, six clonal replicates were grown in 5mL of LB+Cam broth at 30°C in a static incubator for 6hrs, and then washed in minimal media (M9 + Glucose (0.4%) 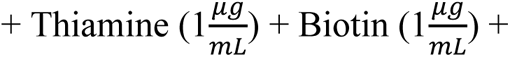 chloramphenicol 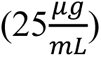). Cultures were diluted with minimal media to an OD600=0.1 using formula provided in *Drosophila* Gut Microbiome Interaction methods. Cultures were further diluted 1:100 in both minimal media and minimal media supplemented with lysine (0.0134%) or isoleucine (0.0065%), for ΔLys or ΔIle, respectively. A sterile assembled BioMe plate was loaded with diluted cultures in appropriate wells – for each pore size, n=4 for amino acid diffusion assay, n=1 for negative control with no supplemental amino acid, and n=1 for positive control with auxotroph loaded with supplemental amino acid in same well. BioMe plate was run on the plate-reader for 96hrs, 30°C, with OD600 measurements at 15mins intervals.

Similar growth, wash, and dilution procedure was followed for the syntrophic co-culture experiment. For each pore size, n=3 for co-culture assay across membrane, n=1 for both negative controls without co-culture partner, and n=1 for positive control with co-culture partners in same well. Plate-reader was run with BioMe plate for 96hrs, 30°C, with OD600 measurements at 15mins intervals. At end of kinetic read, a 5μL aliquot from each well was cultured on minimal media agar plates supplemented with either lysine or isoleucine to determine instances of cross-over. Plates were cultured for 72hrs at 30°C in a static incubator.

### Computational Modeling

We developed a computational model to simulate the interaction between *E. coli* amino acid auxotrophs in the BioMe device. Metabolite diffusion is modeled by a gradient driven flux across the porous membrane. Metabolite uptake flux is bounded by Michaelis-Menten kinetic uptake equations, and growth rate is determined by the minimum biomass flux that can be produced based on metabolite uptake rates and biomass stoichiometry. Amino acid leakage is determined by a stoichiometric parameter specifying the amount of amino acid leaked into the environment for each gram of *E. coli* biomass produced. The uptake, growth constraints, and leakage components of this model are analogous to previously developed dynamic flux balance analysis models (11). Thus our model is essentially a simplified dynamic flux balance analysis model with a growth rate proportional leakage term and gradient driven diffusion between compartments. The full model dynamics are presented in supplemental Equations S2 and the code is available at https://github.com/segrelab/co_culture_device. Simulation of the model was implemented in MATLAB. Testing of run-time and accuracy was conducted to benchmark three MATLAB ODE solvers (ode23tb, ode45, and ode15s). The function ode23tb was found to have the best performance, with consistent accuracy when using maximum step sizes of 0.0025 hrs. Thus, this solver function and maximum step size setting was used to simulate model dynamics.

We implemented an approximate Bayesian computation based approach to integrate experimental data with computational modeling. Approximate Bayesian computation is used to infer the posterior distribution of the parameters of a computational model by using simulations of experimental results to approximate the likelihood of the data (60, 61). We implemented a rejection based algorithm where sample parameters are initially drawn from a given prior distribution and are included in the posterior distribution if the difference between the simulated and experimental value of a given statistic is less than some specified threshold. The statistic that we utilized was the log base 10 of the ratio between growth at 48 hours of the *E. coli* auxotrophs in the same well (positive control) divided by the growth in opposite wells (membrane-separated) (Supplemental Equations 4). Using this ratio allowed us to compare experimental results to simulated results without calibrating between biomass units, as our optical density growth measurements were made ratio-scale by subtracting a blank control. The threshold that we used was two standard deviations from the experimentally measured mean ratio.

We began by sampling diffusion and leakage parameters while assuming that the leakage of isoleucine and lysine were equal (Figure 7). The uptake kinetics and biomass parameters were estimated from the literature and were fixed at their estimated values (prior distribution was a Dirac delta function) (Supplemental Equations S3). The leakage and diffusion parameters were randomly sampled from a log uniform prior distribution ranging from 10^-6^ to 10^-3^ L/hr for the diffusion rate and 10^-2^ to 10 mmol/g for the leakage stoichiometry. Next, we used our model to investigate the unequal growth of the ΔLys and ΔIle auxotrophs (Figure S7.1). We fixed the diffusion rate to a value on the high-end of what we observed for the diffusion of the chemical dyes (3.68*10^-5^ L/hr). Then, sampled the leakage for ΔLys and ΔIle independently from log uniform prior distributions with the same ranges as used previously. Finally, we added noise to all of the literature estimated parameters (with the exception of volume) to represent an increased level of uncertainty (Figure S7.2 and S7.3). We sampled all literature estimated parameters from a log uniform prior distribution that varied by 1 order of magnitude around the originally fixed literature estimated value. This more uncertain prior distribution was used to repeat the inference of leakage stoichiometry for both the equal (Figure S7.2) and unequal (Figure S7.3) cases.

### Data Availability

All data and code are available on GitHub at https://github.com/segrelab/co_culture_device. Data includes raw kinetic optical density measurements for the small molecule dye diffusion, *Drosophila* gut microbe co-culture, amino acid diffusion, and syntrophic *E. coli* auxotroph co-culture experiments, and MATLAB scripts for generation of all figures from raw data are provided in the “raw_data_and_plots” subdirectory. MATLAB scripts for solving the differential equations of the computational model and for the Bayesian analyses are located in the “modeling” subdirectory.

## Acknowledgements

We are grateful to members of the Segrè lab for helpful feedback and constructive advice, and to Harris Wang and Ali Zomorrodi for making the *E. coli* auxotroph strains available to us. Specifically, we would like to thank Melisa Osborne for assistance with experimental protocols and training. We would also like to acknowledge the staff of the Boston University Engineering Product Innovation Center and Jonathan Perry of the Biology Department Workshop for assistance in the initial fabrication of the device. This work was partially supported by grants from the National Institutes of Health (National Institute of General Medical Sciences, award R01GM121950; National Institute of Dental and Craniofacial Research, award number R01DE024468), the National Science Foundation (grants 1457695 and NSFOCE-BSF 1635070), the Human Frontiers Science Program (RGP0020/2016), and the Boston University Interdisciplinary Biomedical Research Office. DBB acknowledges support by the Boston University training program in quantitative biology and physiology under Ruth L Kirschstein National Research Service Award T32GM008764 from the National Institute of General Medical Sciences.

## Supplement

**Supplemental Table S1:**
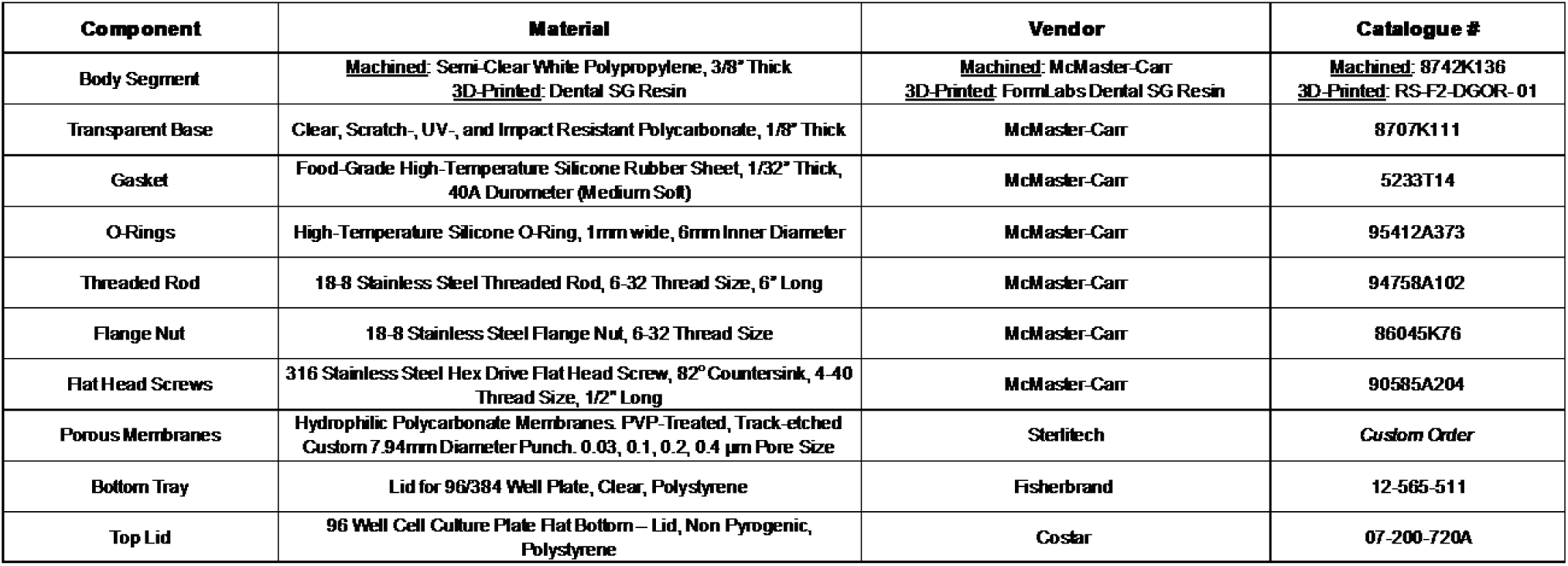
BioMe Materials.

**Supplemental Figure S2.1:**
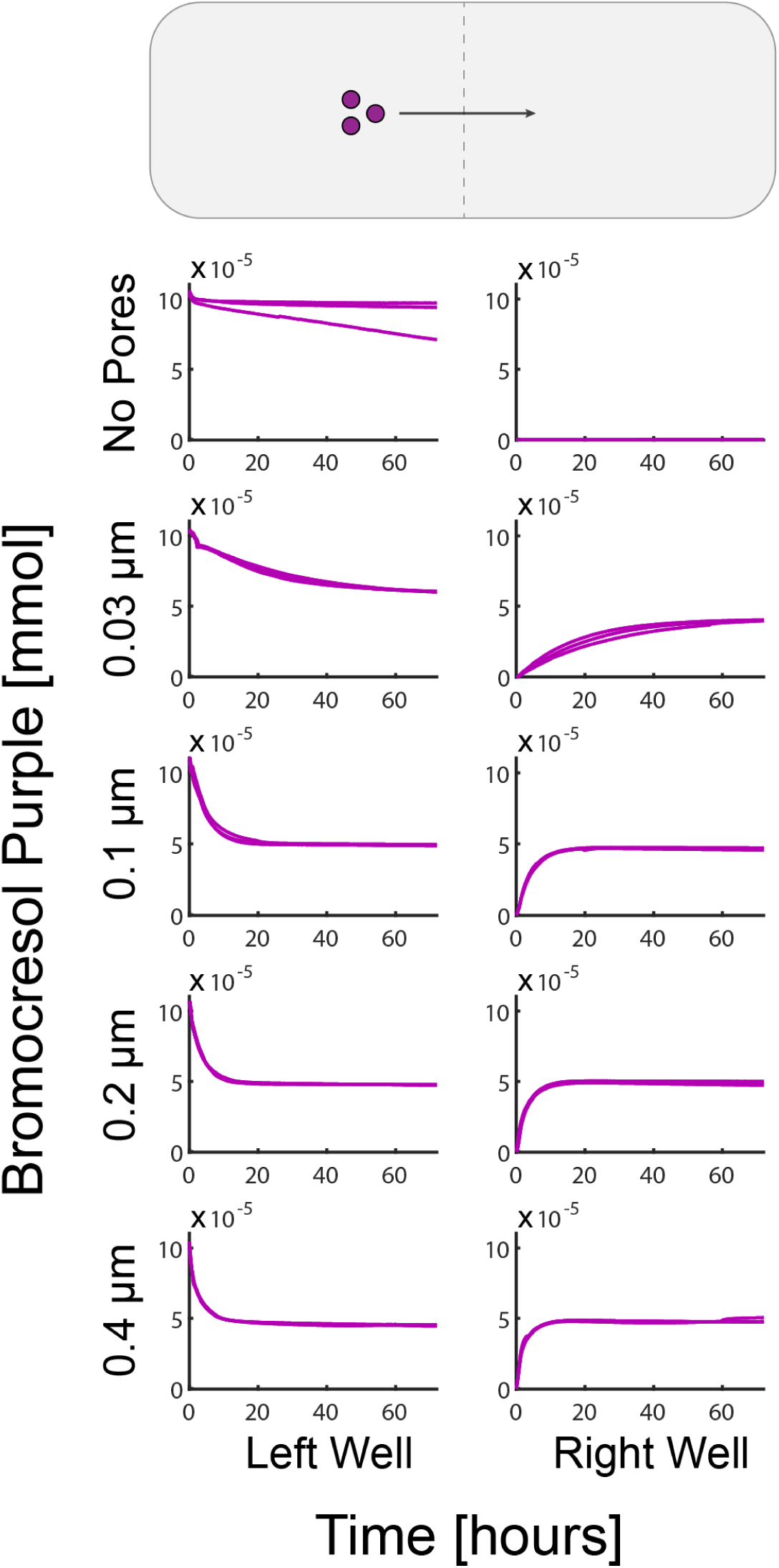
Bromocresol Purple Diffusion Across the BioMe Plate. Small-molecule dye (bromocresol purple) was placed on one side of the membrane in the BioMe plate for membranes of different pore sizes. Three replicates are shown here (overlapping) for each pore size. The amount of dye was measured by optical density at the isosbestic point (optical density was converted to amount by a linear calibration shown in supplemental Figure S2.2).

**Supplemental Figure S2.2:**
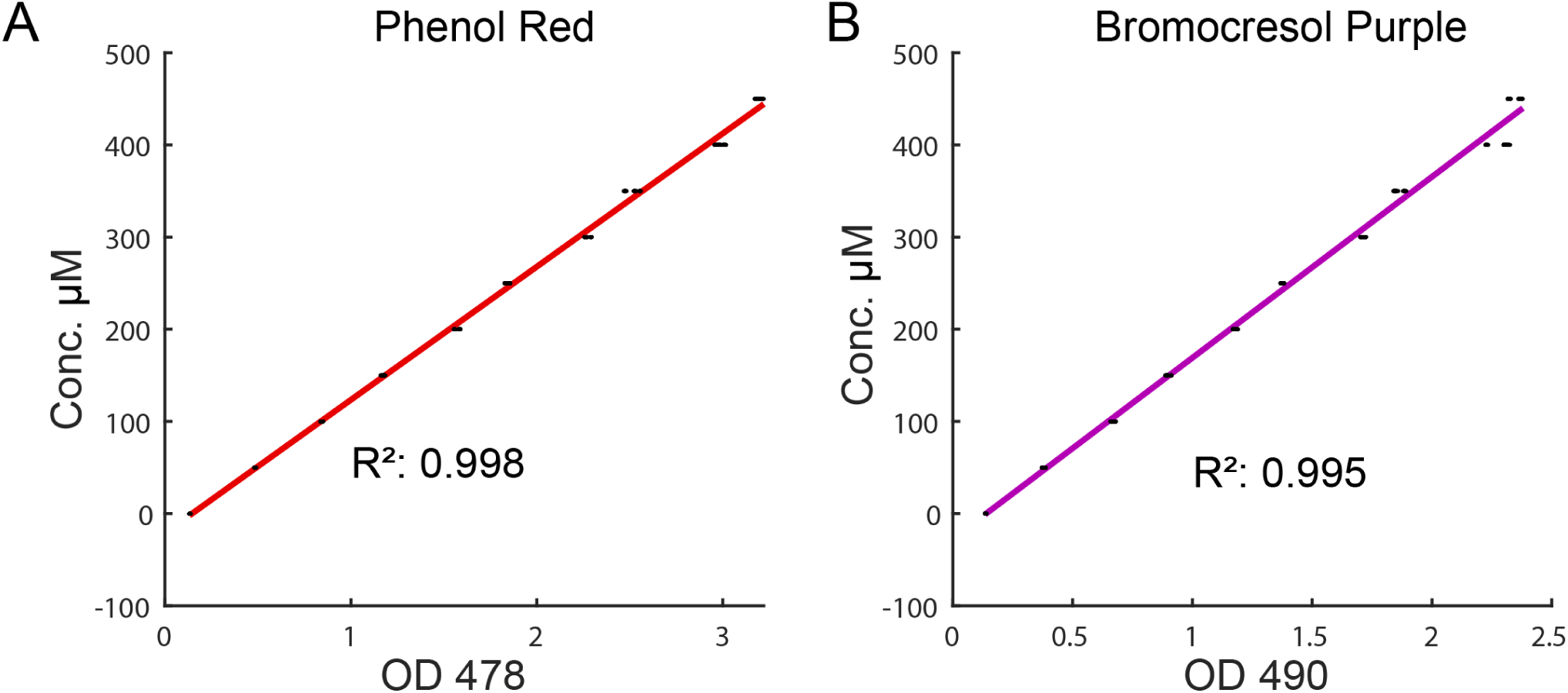
Small Molecule Dye Calibration Curves. A linear calibration converting phenol red **(A)** and bromocresol purple **(B)** optical density at the isosbestic point to concentration.

**Supplemental Figure S2.3:**
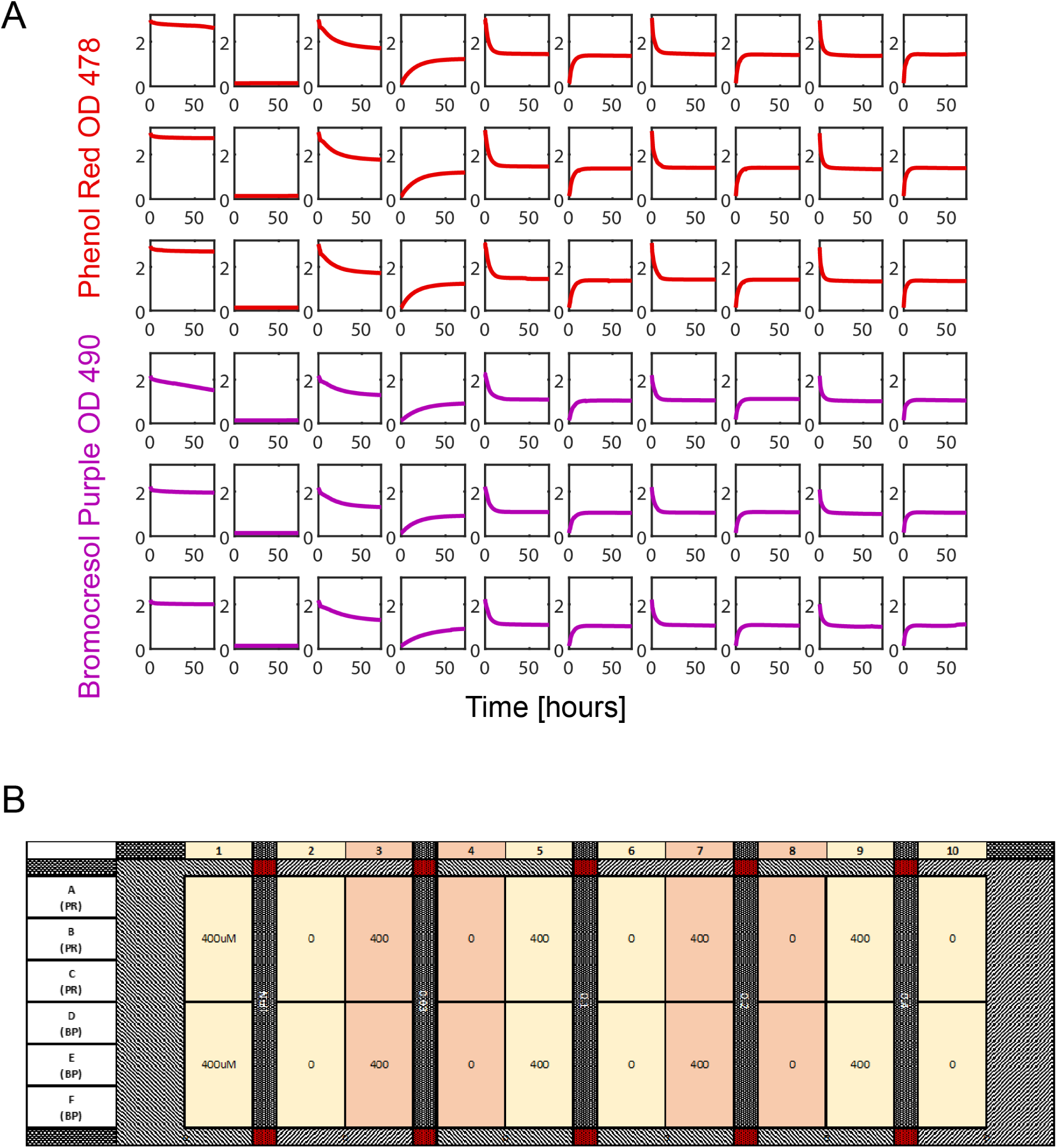
Dye Diffusion Plate layout. **(A)** The raw optical density readings from the diffusion experiment are shown. Red – phenol red, Purple – bromocresol purple**. (B)** The layout of the experiment on the BioMe plate is shown.

**Supplemental Figure S2.4:**
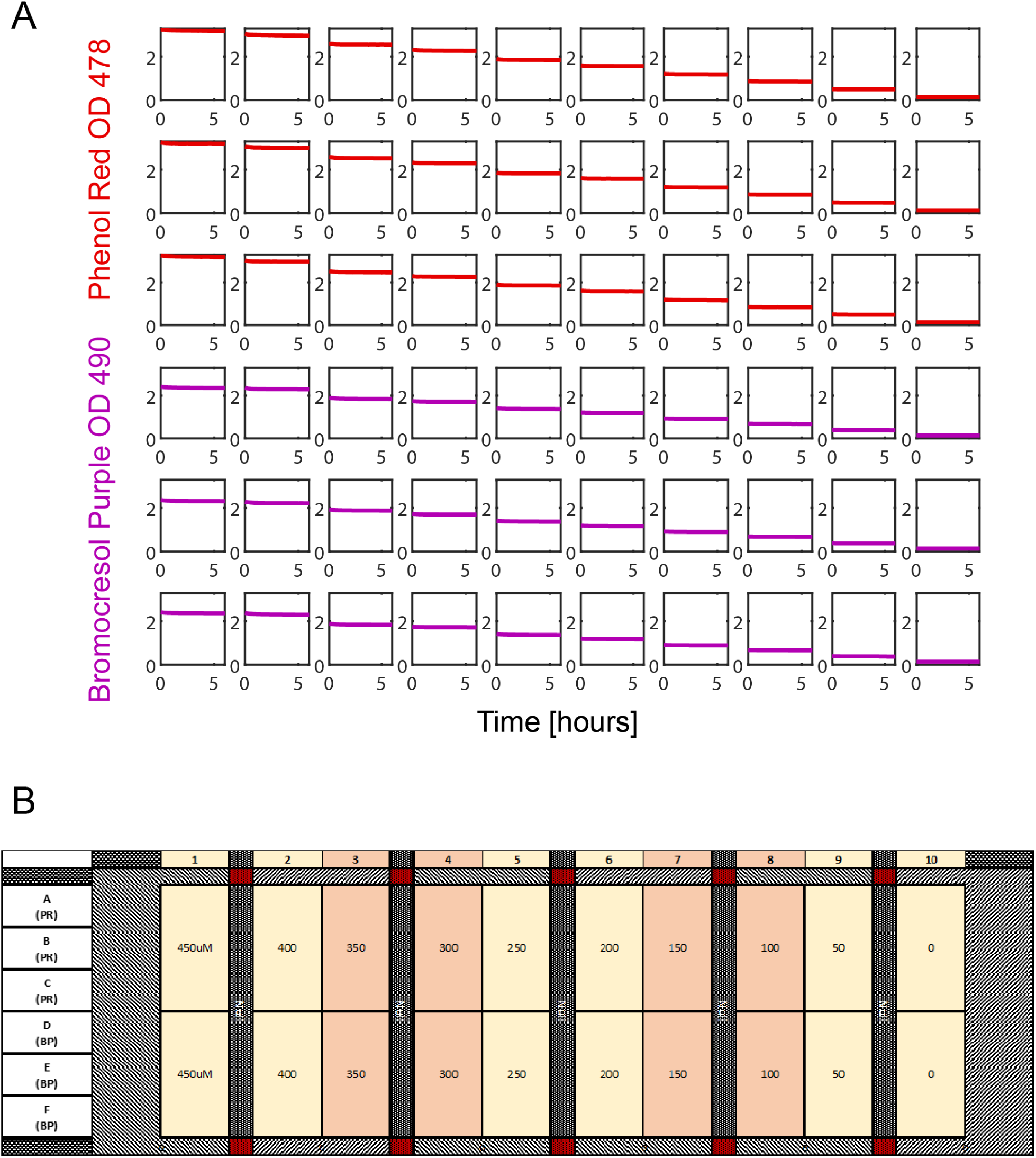
Dye Calibration Plate layout. **(A)** The raw optical density readings from the dye calibration are shown. Red – phenol red, Purple – bromocresol purple**. (B)** The layout of the experiment on the BioMe plate is shown.

**Supplemental Equations S1:**
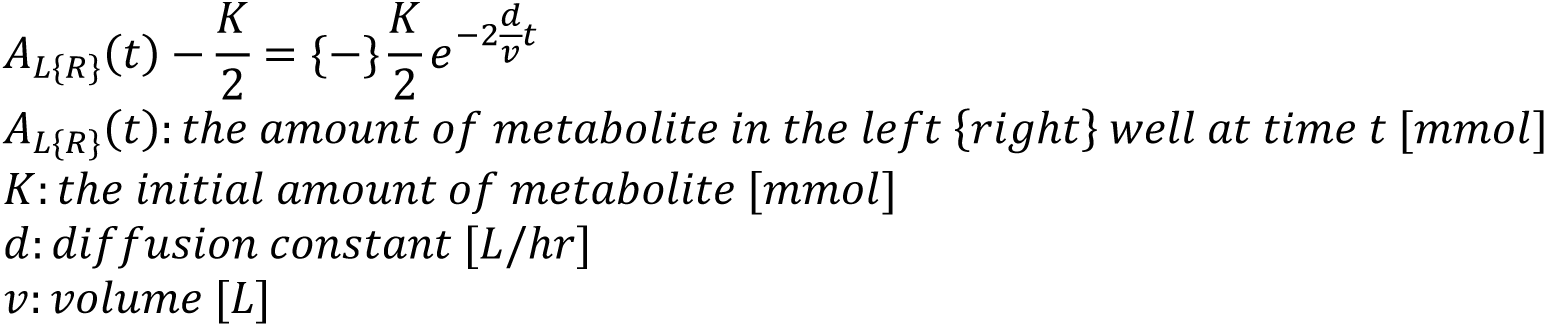
Metabolite diffusion dynamics.

**Supplemental Figure S2.5:**
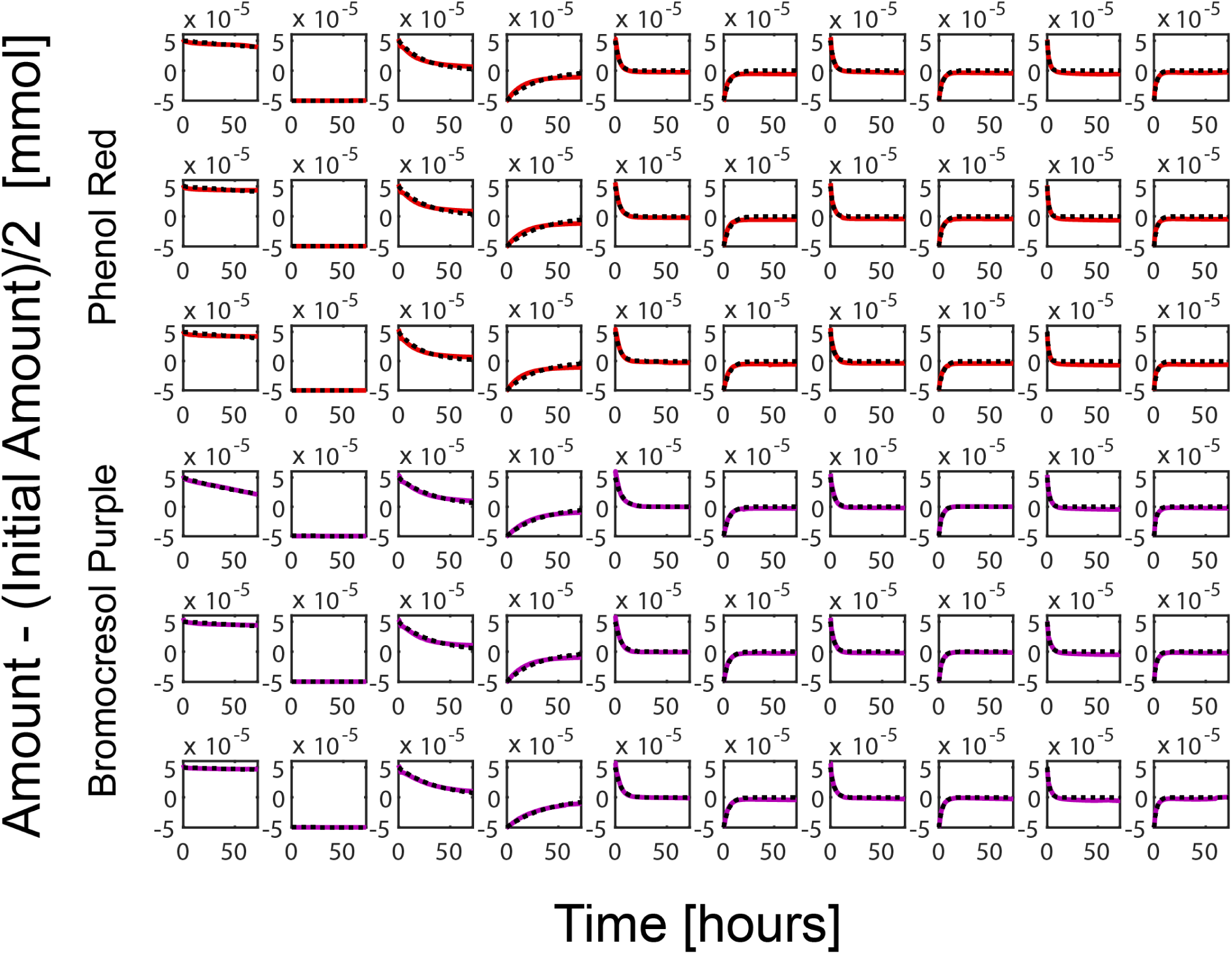
Metabolite diffusion curves fit to exponential function. The experimental data from the chemical dye diffusion experiment was used to plot the value on the left hand side of supplemental Equations S1 (red – phenol red, purple – bromocresol purple). An exponential function was then fit to this data using the form of the right hand side of supplemental Equations S1 (dotted black line).

**Supplemental Figure S3.1:**
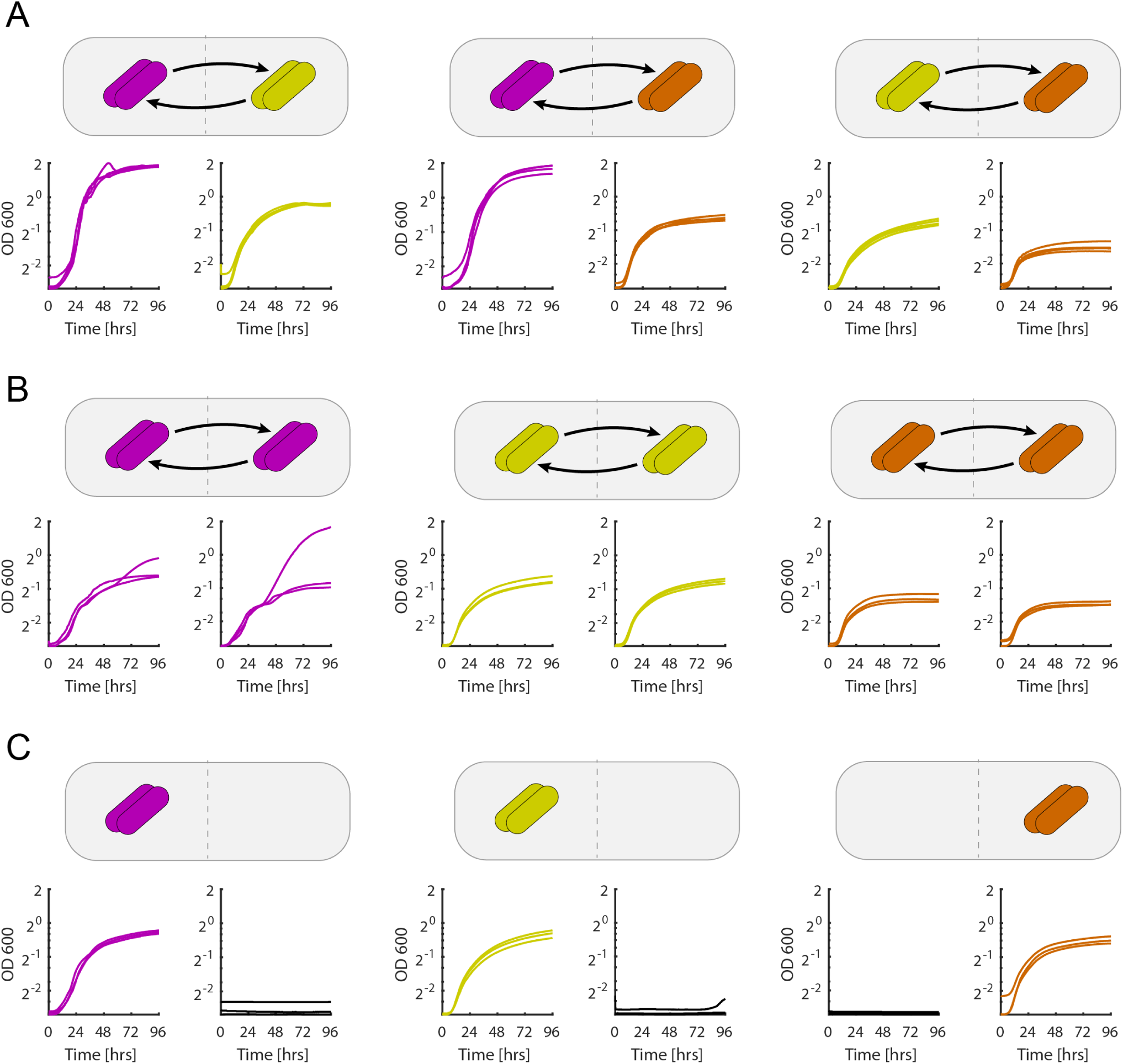
*Drosophila* gut microbe interactions - all data and replicates. Growth curves and replicates for all co-culture experiments between *A. oryzifermentans* (purple), *L. plantarum* (yellow), and *L. brevis* (orange) are shown. The schematic demonstrates the type of experiment performed: **(A)** co-culture, **(B)** self-interaction (self-control), and **(C)** empty well interaction (media-control).

**Supplemental Figure S3.2:**
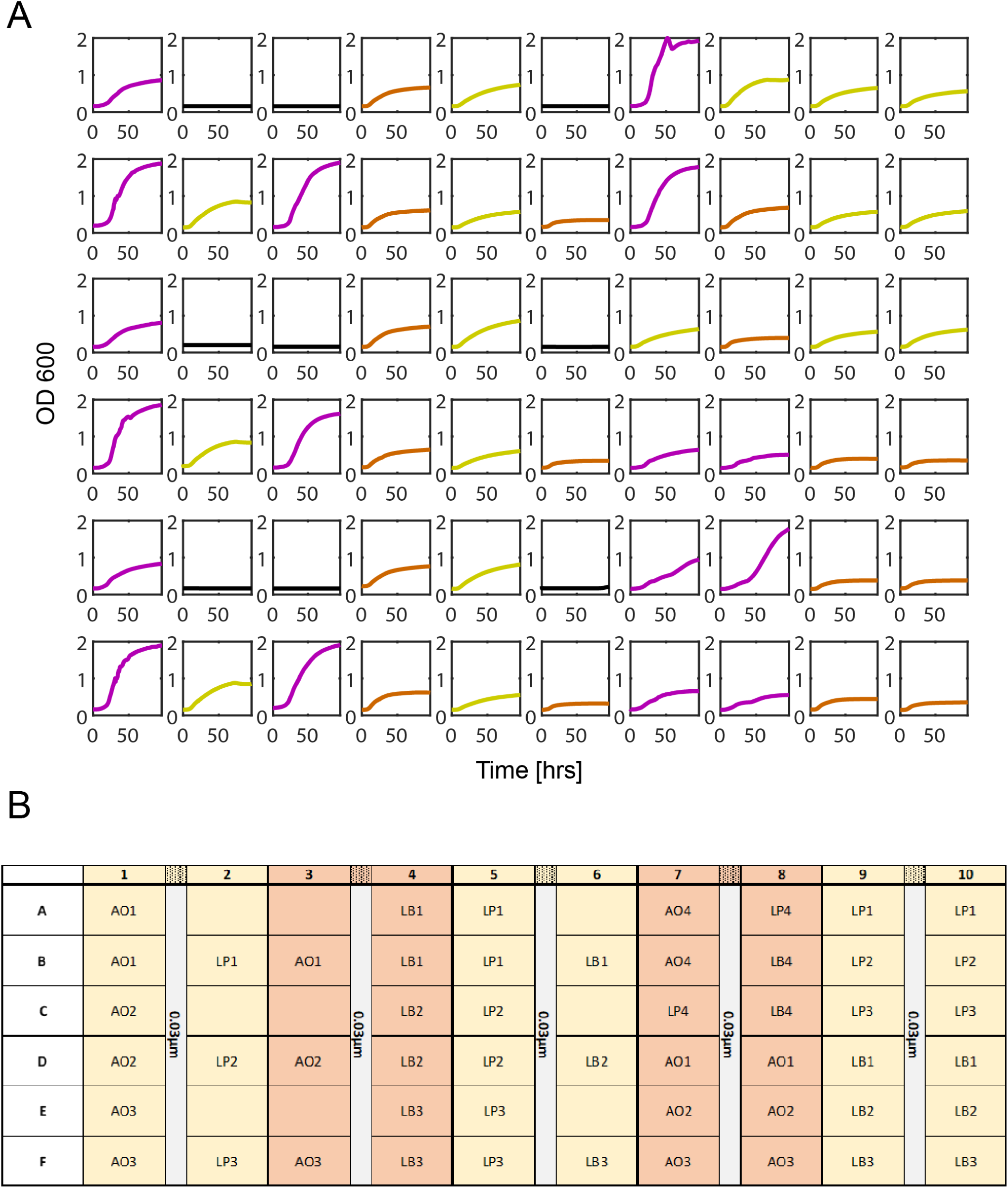
*Drosophila* gut microbe interactions - plate layout. **(A)** The raw experimental optical density measurements for all cultures are shown for co-culture experiments of *A. oryzifermentans* (purple), *L. plantarum* (yellow), and *L. brevis* (orange). **(B)** The layout of the experiment on the BioMe plate is shown. Two outliers for the *A. oryzifermentans* v.s. self-co-cultures were left out of the analysis done in figure 3, these are the curves from the interacting wells E7 and E8.

**Supplemental Figure S4.1:**
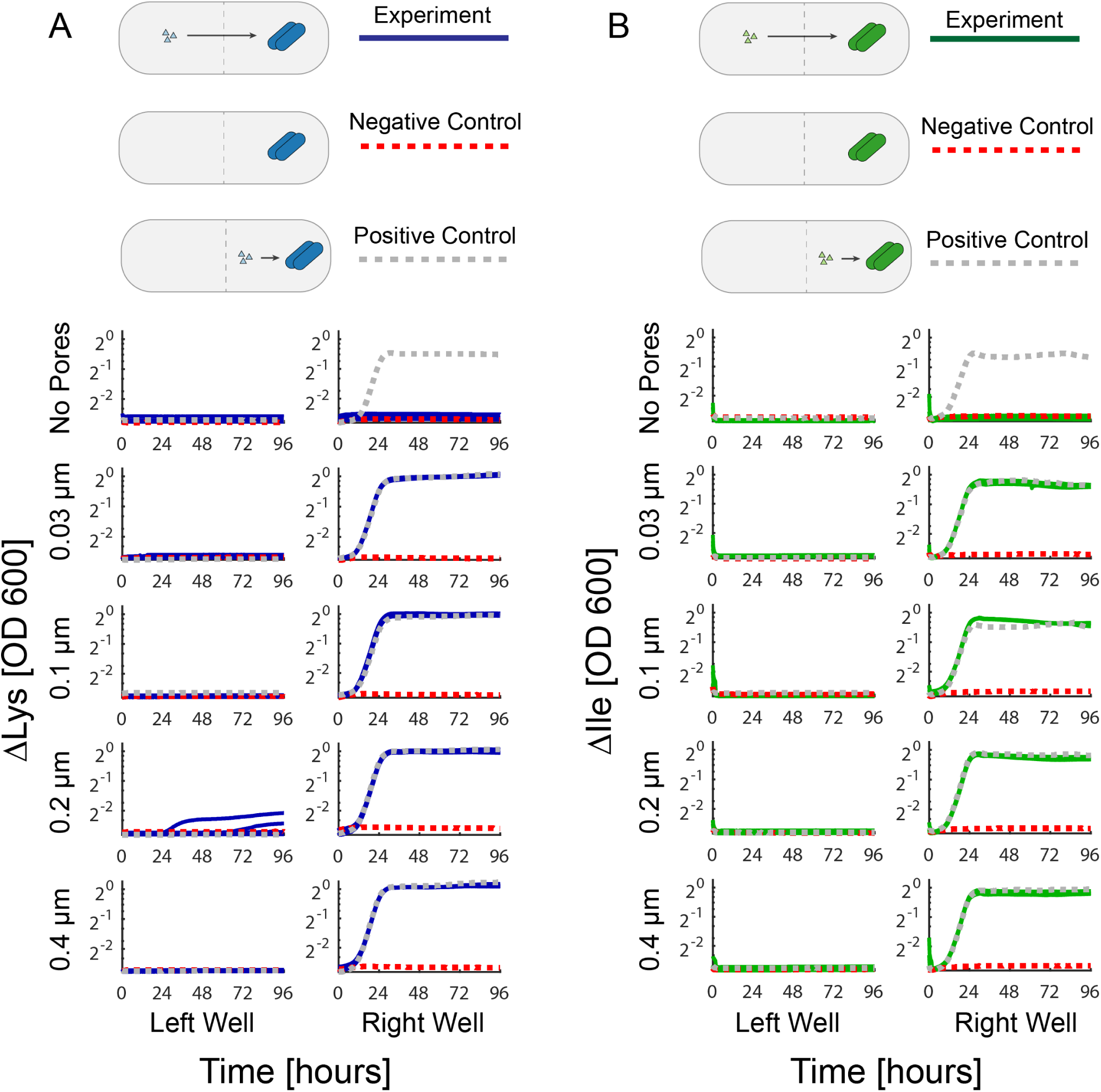
*E. coli* auxotroph amino acid diffusion experiments - all data and replicates. The growth of ΔLys **(A)** and ΔIle **(B)** are shown for five different pore sizes (No Pores, 0.03, 0.1, 0.2, and 0.4 μm) alongside positive and negative control growth curves. Blue – ΔLys, Green – ΔIle, Red – negative controls, Gray – positive controls.

**Supplemental Figure S4.2:**
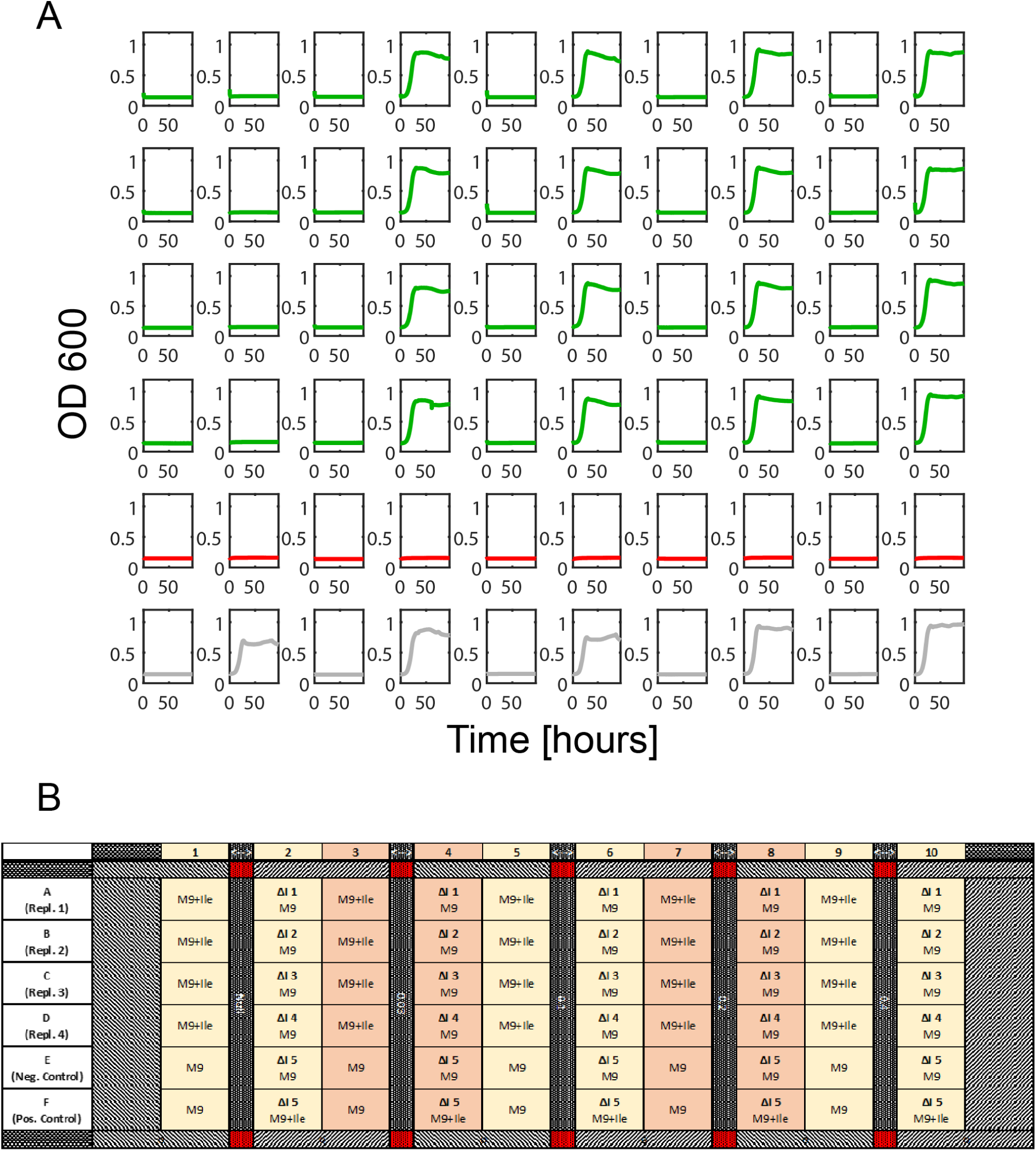
*E. coli* ΔIle auxotroph amino acid diffusion experiments - plate layout. **(A)** The raw optical density readings for the *E. coli* ΔIle auxotroph isoleucine diffusion experiment are shown. Green – ΔIle, Red – negative controls, Gray – positive controls. **(B)** The layout of the experiment on the BioMe plate is shown.

**Supplemental Figure S4.3:**
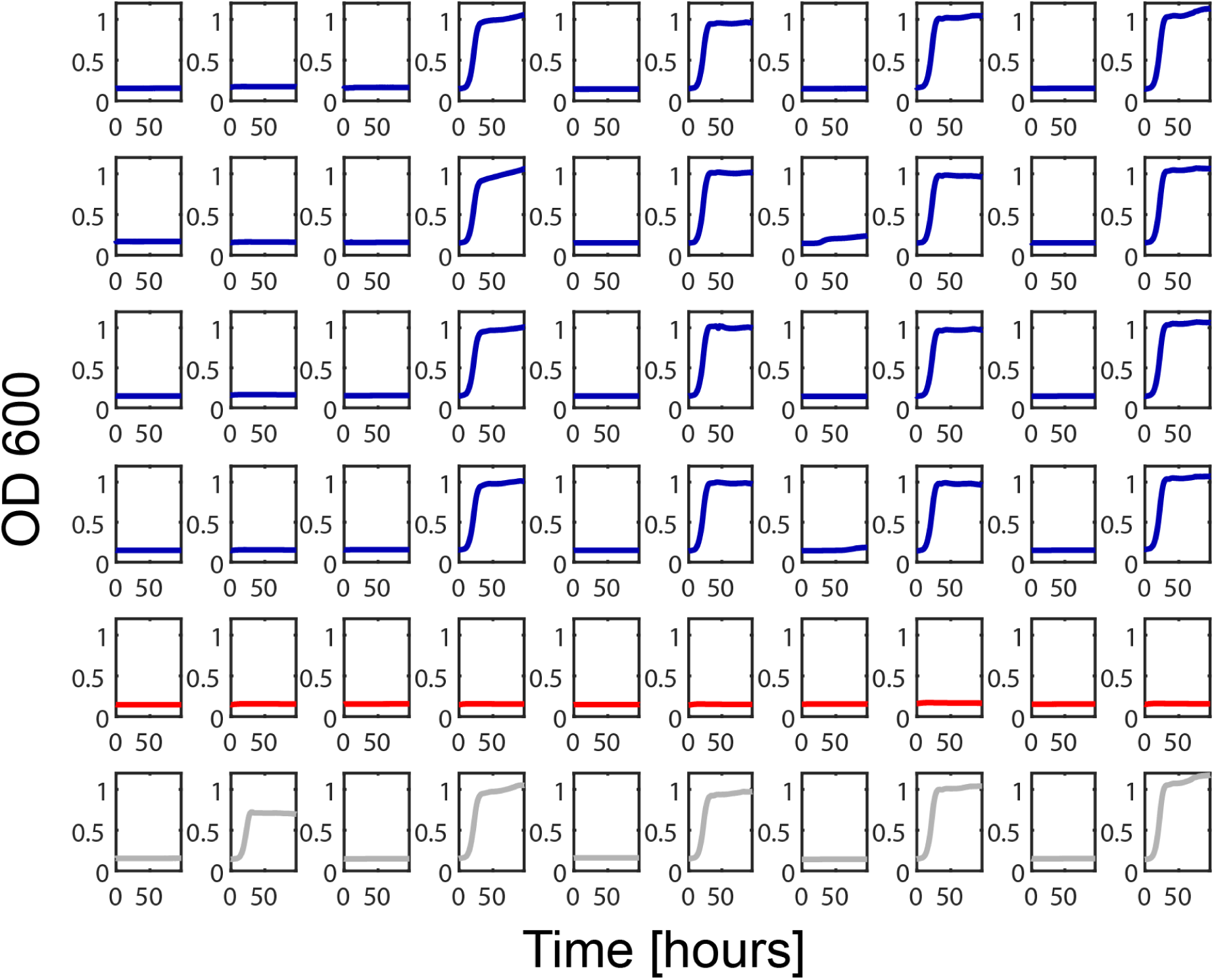
*E. coli* ΔLys auxotroph amino acid diffusion experiments - plate layout. **(A)** The raw optical density readings for the *E. coli* ΔLys auxotroph lysine diffusion experiment are shown. The layout of the experiment on the BioMe plate is the same as that shown for the ΔIle experiment, with ΔLys replacing ΔIle and lysine replacing isoleucine. Blue – ΔLys, Red – negative controls, Gray – positive controls.

**Supplemental Figure S4.4:**
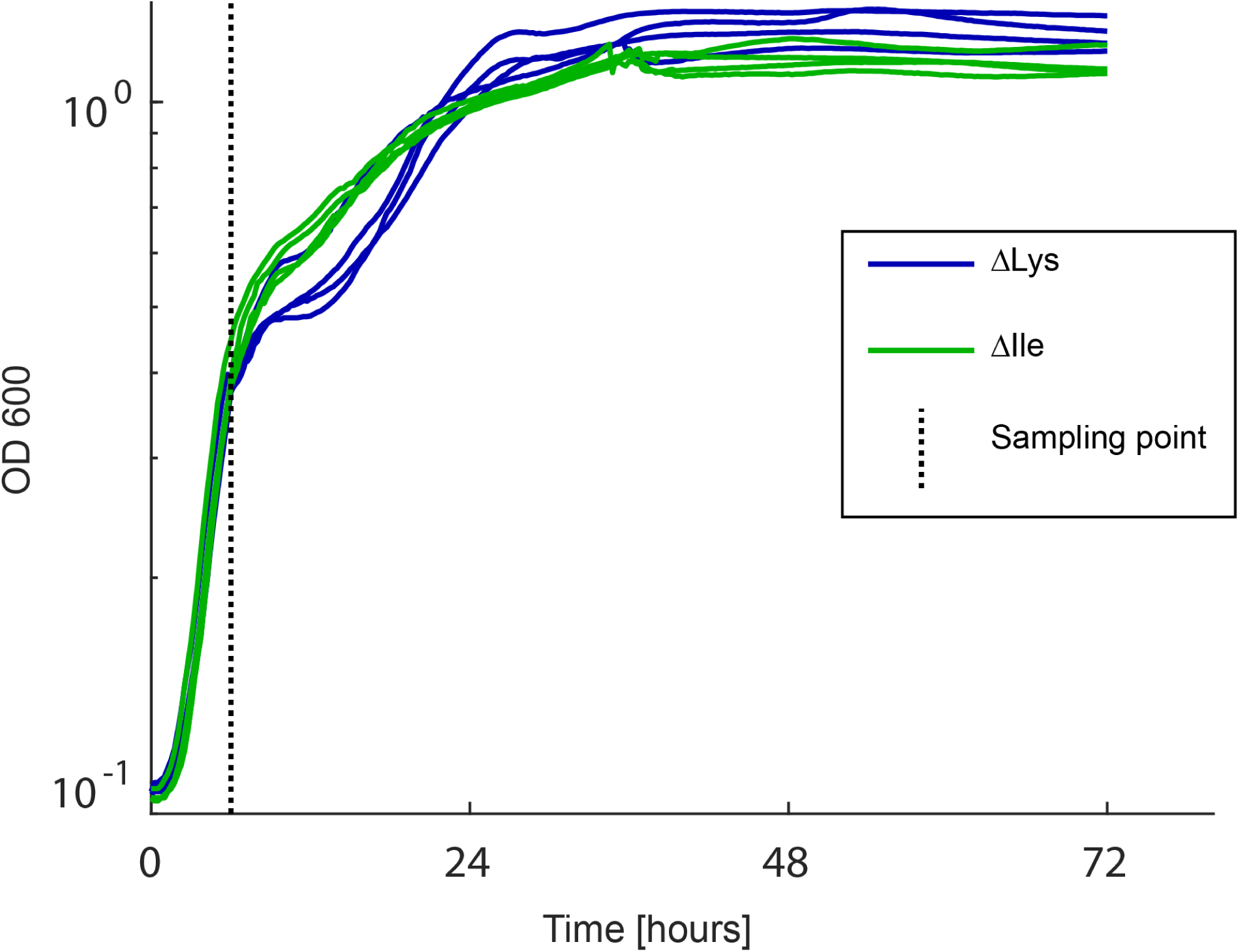
Initial growth curves for ΔLys and ΔIle in LB medium. Initial cultures for experiments were diluted from the sampling point shown by the dotted black line. Four replicates for both ΔLys and ΔIle.

**Supplemental Figure S5.1:**
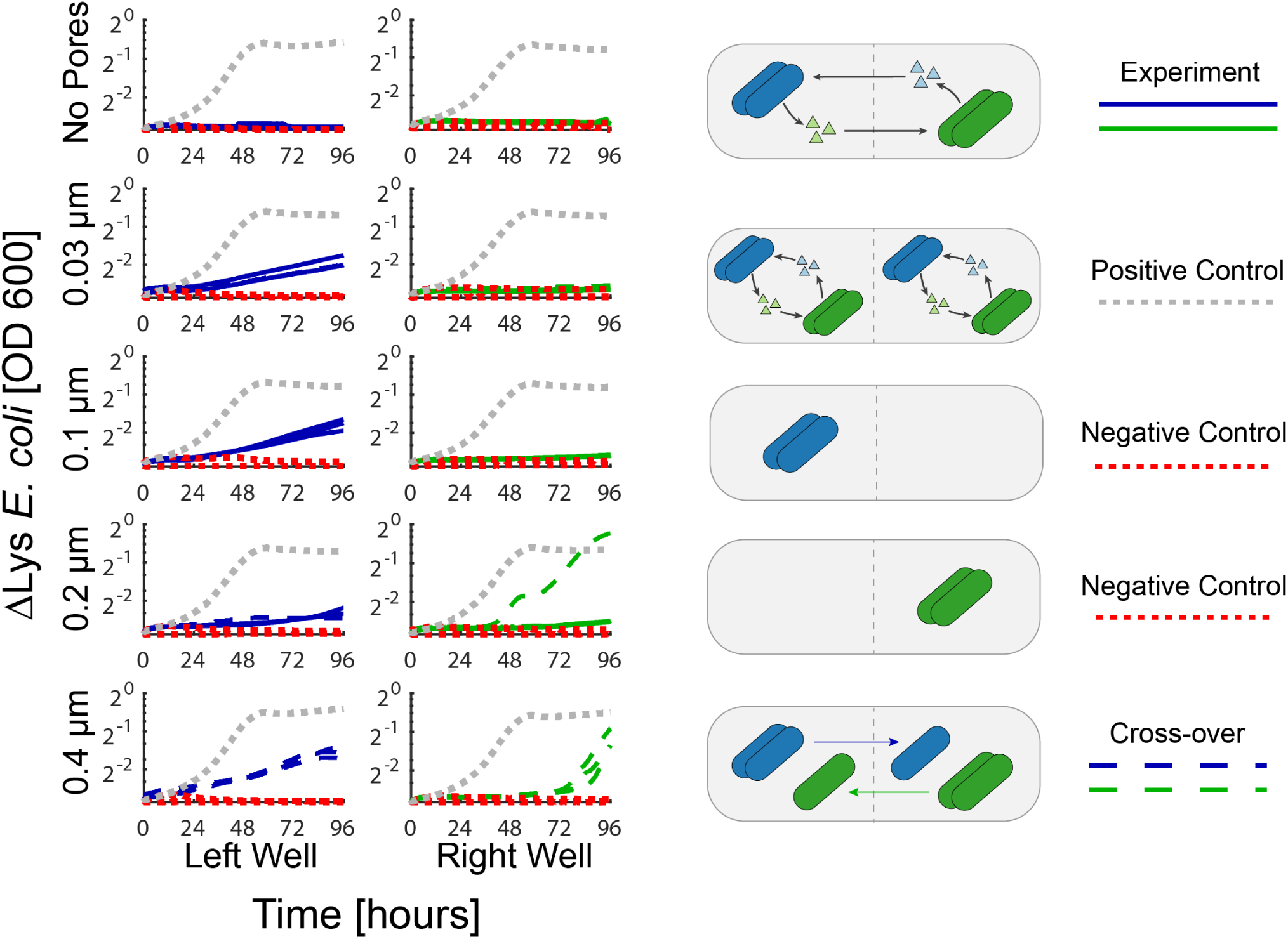
*E. coli* auxotroph co-culture experiments - all data and replicates. The growth curves of ΔLys and ΔIle co-cultures are shown for 5 different pore sizes (No Pores, 0.03, 0.1, 0.2, and 0.4 μm) alongside positive and negative controls. Dotted lines indicate cross-over of auxotrophs as indicated in supplemental Table S3. Blue – ΔLys, Green – ΔIle, Red – negative controls, Gray – positive controls.

**Supplemental Figure S5.2:**
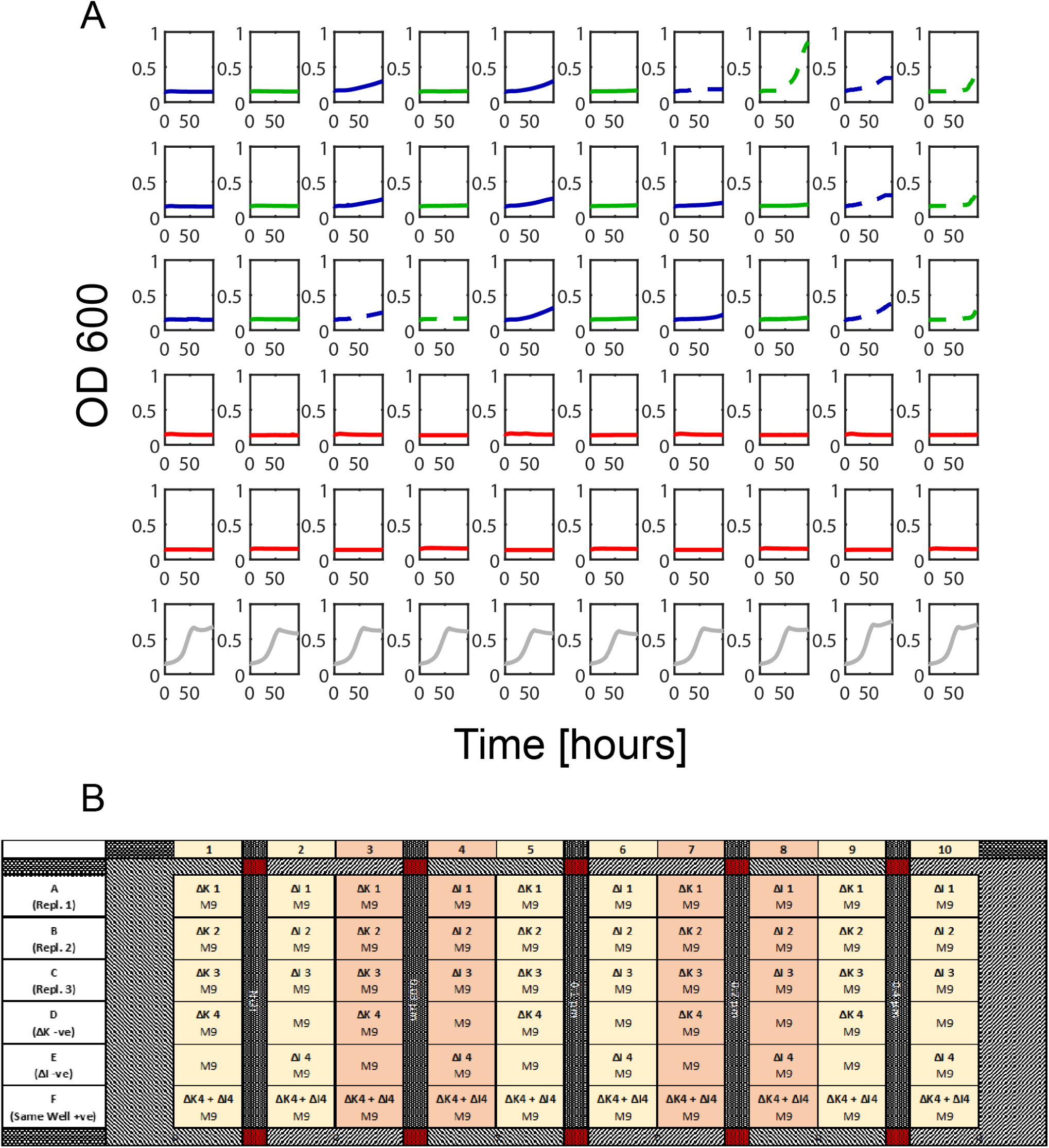
*E. coli* auxotroph co-culture experiments - plate layout. **(A)** The raw optical density measurements for the *E. coli* auxotroph co-culture experiment are shown. Dotted lines indicate cross-over of auxotrophs as indicated in supplemental Table S3. Blue – ΔLys, Green ΔIle, Red – negative controls, Gray – positive controls. **(B)** The layout of the experiment on the BioMe plate is shown.

**Supplemental Table S2:** Cross-over assay. A selective platting assay at the end of the *E. coli* auxotroph co-culture experiment was performed to identify when organisms had crossed into the adjacent well of the BioMe plate. Black letters indicate that the correct auxotroph was identified in that well, and red letters indicate that the incorrect auxotroph was found in the well.

|  | 1 |  | 2 | 3 |  | 4 | 5 |  | 6 | 7 |  | 8 | 9 |  | 10 |
| --- | --- | --- | --- | --- | --- | --- | --- | --- | --- | --- | --- | --- | --- | --- | --- |
| A<br>(Repl. 1) | ΔK | No Pores | ΔI | ΔK | 0.03 μm | ΔI | ΔK | 0.1 μm | ΔI | ΔK | 0.2 μm | ΔK & ΔI | ΔK | 0.4 μm | ΔK & ΔI |
| B<br>(Repl. 2) | ΔK |  | ΔI | ΔK |  | ΔI | ΔK |  | ΔI | ΔK |  | ΔI | ΔK |  | ΔK & ΔI |
| C<br>(Repl. 3) | ΔK |  | ΔI | ΔK & ΔI |  | ΔI | ΔK |  | ΔI | ΔK |  | ΔI | ΔK |  | ΔK & ΔI |
| D<br>(Repl. 4) | ΔK |  |  | ΔK |  |  | ΔK |  |  | ΔK |  |  | ΔK |  |  |
| E<br>(-ve Control) |  |  | ΔI |  |  | ΔI |  |  | ΔI |  |  | ΔI |  |  | ΔI |
| F<br>(+ve Control) | ΔK & ΔI |  | ΔK & ΔI | ΔK & ΔI |  | ΔK & ΔI | ΔK & ΔI |  | ΔK & ΔI | ΔK & ΔI |  | ΔK & ΔI | ΔK & ΔI |  | ΔK & ΔI |

**Supplemental Equations S2:**
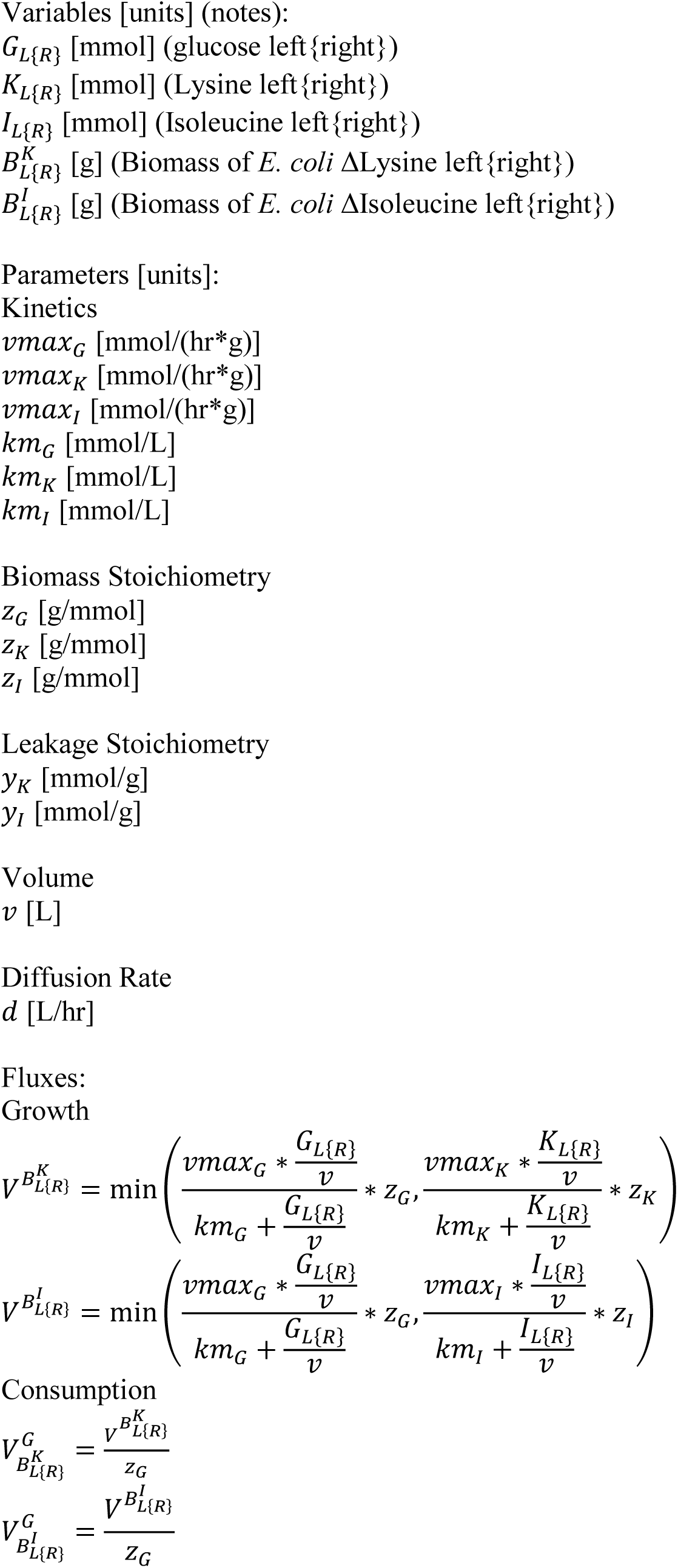
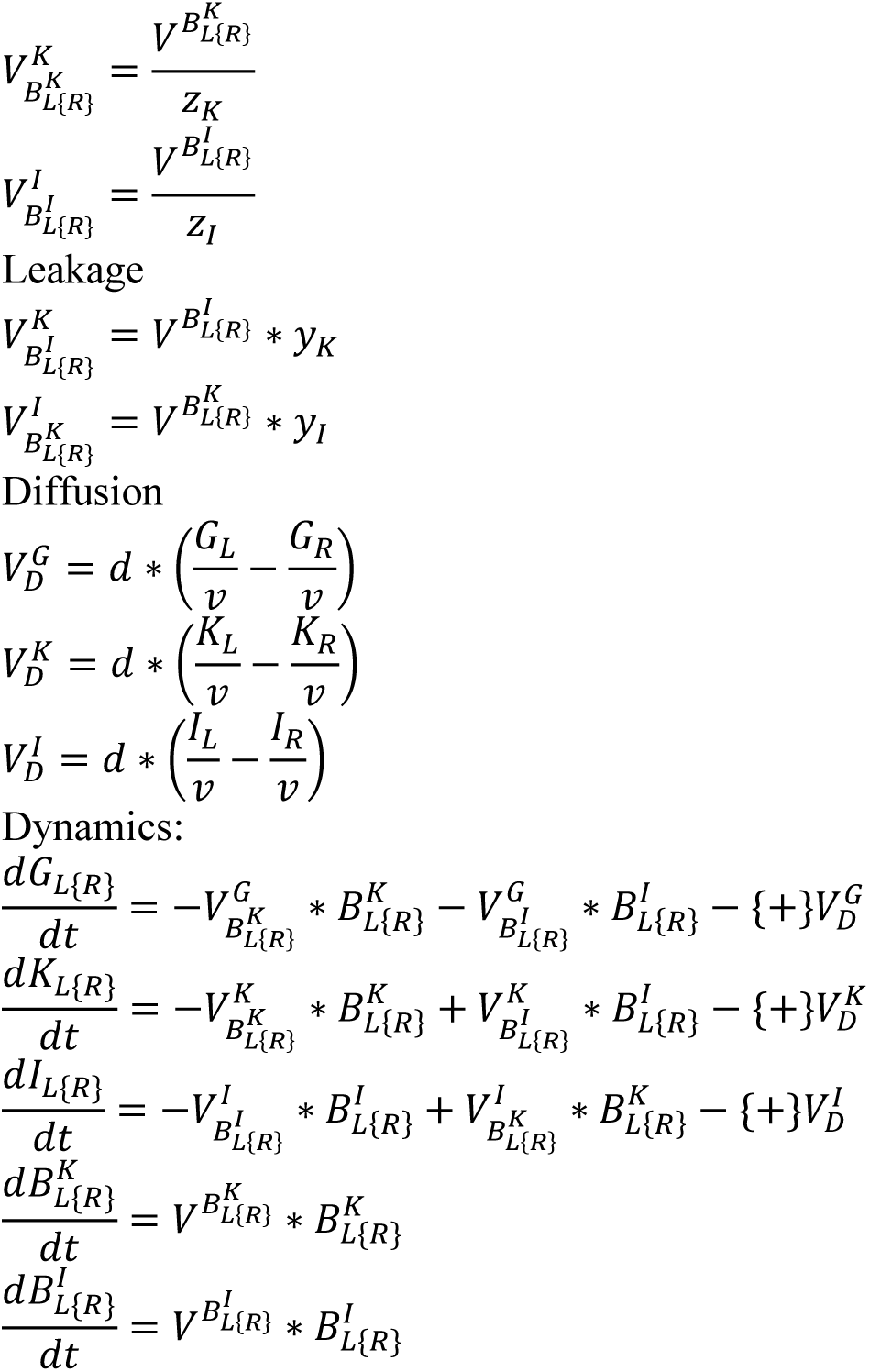
Computational Model of *E. coli* Amino Acid Auxotroph Interaction.

**Supplemental Figure S6.1:**
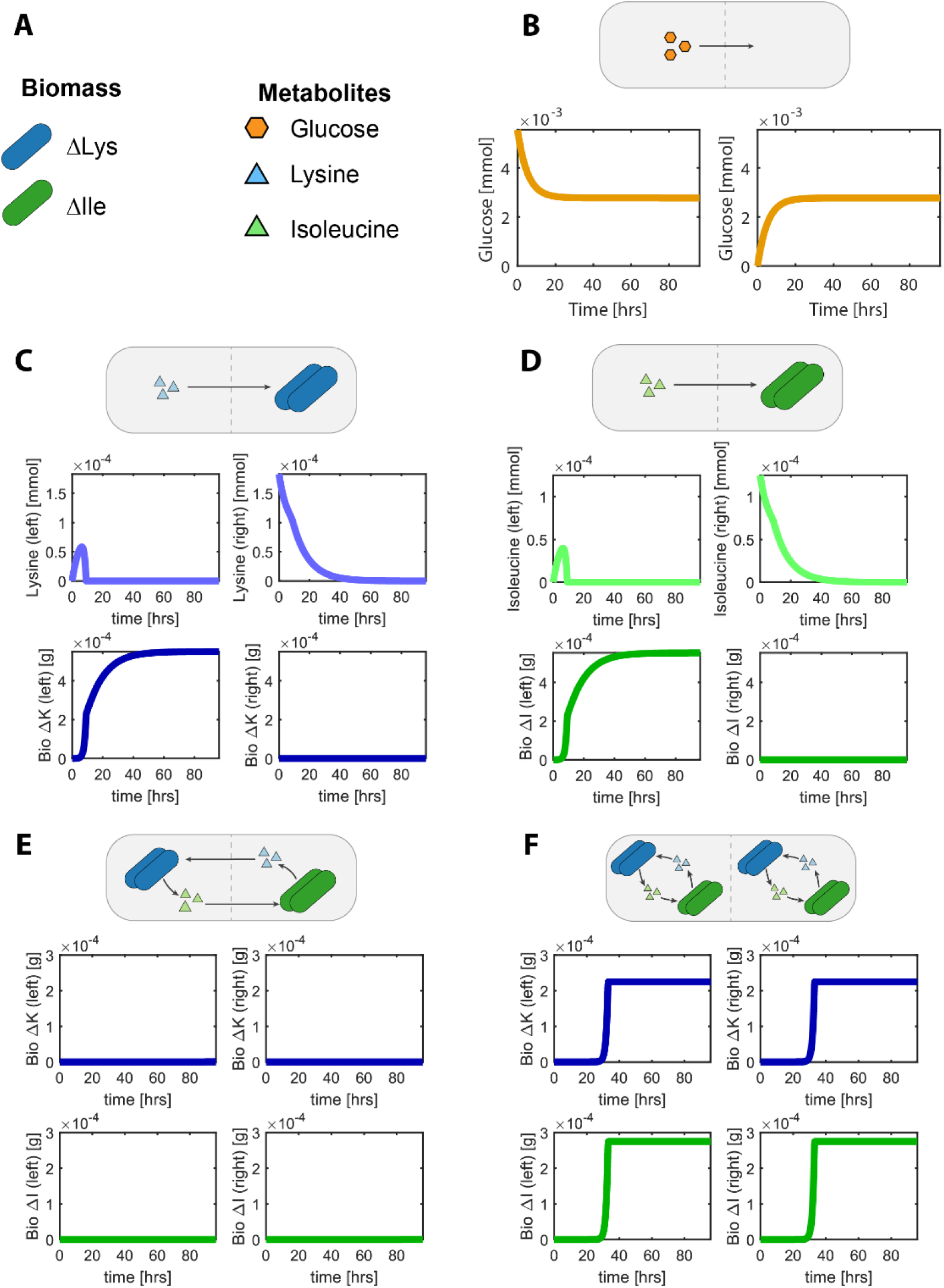
Simulation of *E. coli* amino acid auxotroph experiments. Different experiments can be simulated by changing the initial conditions and parameters of the model. **(A)** The legend describes the variables in the dynamic model. These variables appear in the schematic above each experiment, describing the layout of the experiment performed. **(B)** Chemical diffusion: glucose is added to one side of the device, and diffuses across the membrane until the concentration is equal on both sides. **(C&D)** Amino acid diffusion: Auxotrophic *E. coli* strains are placed on one side of the membrane with their auxotrophy amino acid on the opposite side (C – Lysine, D – Isoleucine). Growth is observed as the amino acid diffuses across the membrane and is taken up by *E. coli.* **(E, F)** Auxotroph co-culture: Auxotrophic *E. coli* strains are placed on the opposite (E) or same (F) sides of the device with no initial amino acid. Growth is observed to be much faster in the same well case (F) than the opposite well case (E).

**Supplemental Equations S3:**
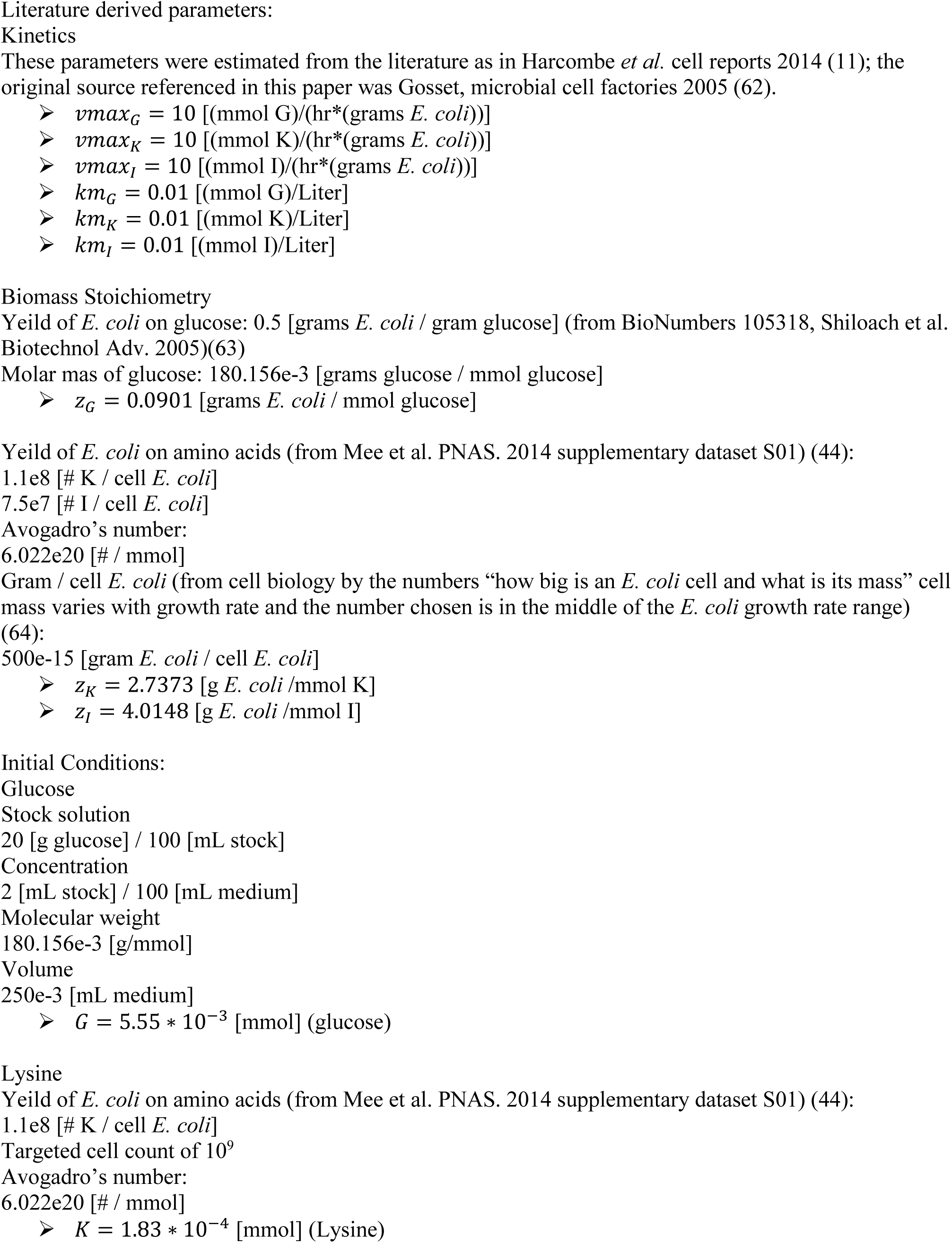
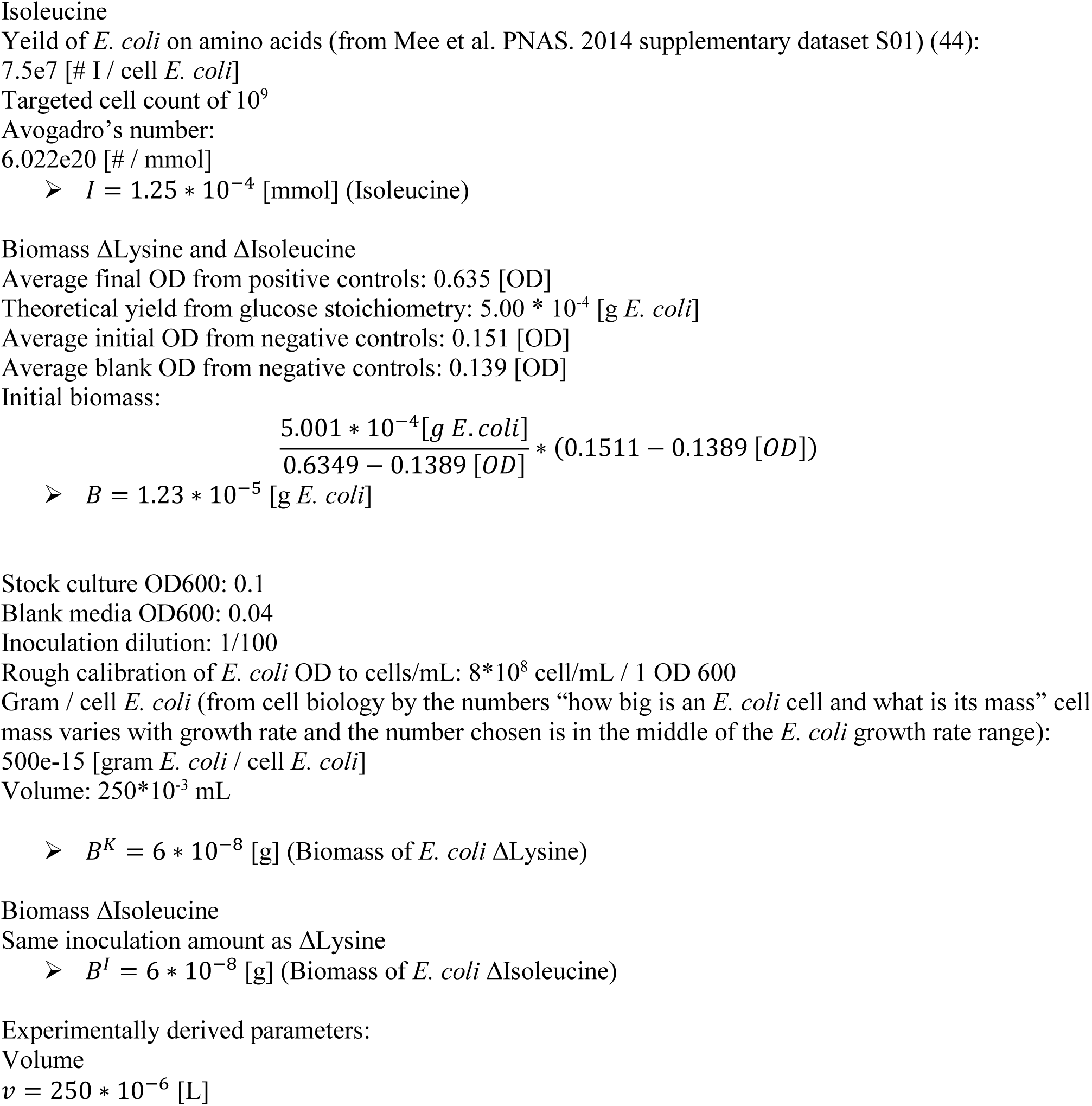
Parameters and Initial Conditions.

**Supplemental Equations S4:**
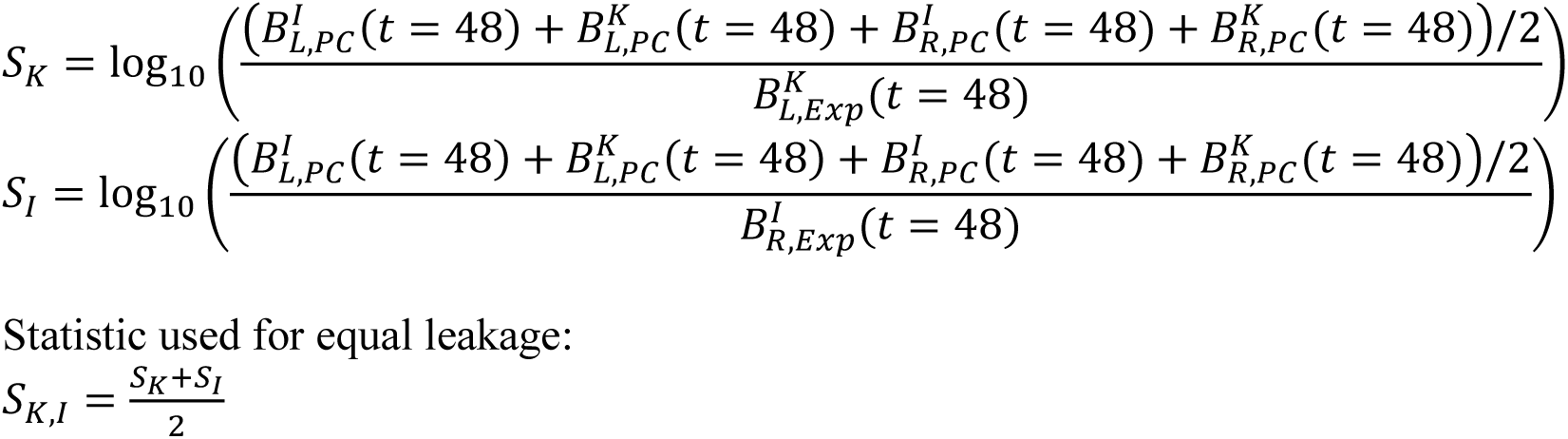
Statistic used for approximate Bayesian computation.

**Supplemental Figure S7.1:**
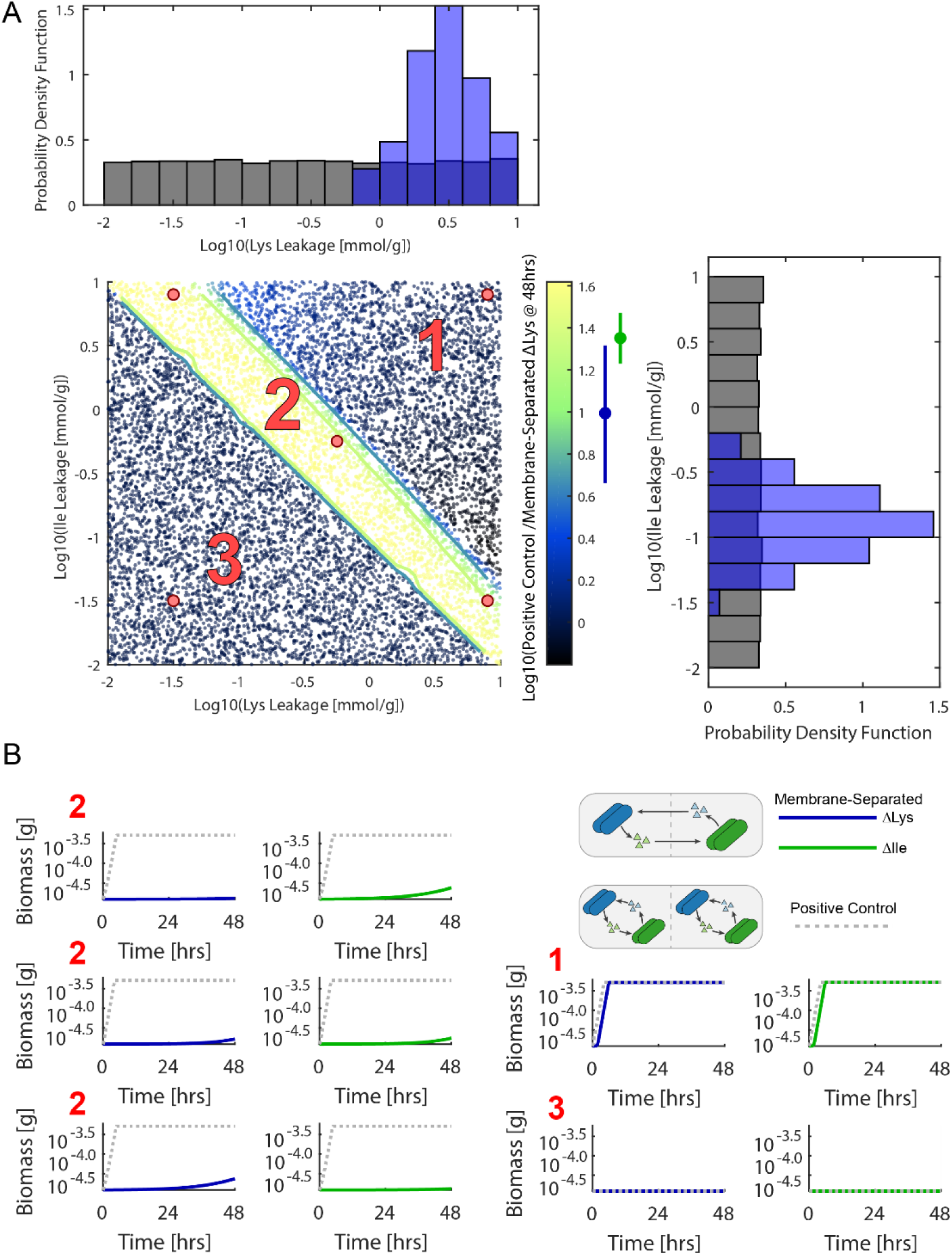
Sampling of unequal leakage parameters. **(A)** Sample space of ΔLys and ΔIle leakage. The statistic used for approximate Bayesian computation, log base 10 of the ratio between positive control growth and membrane-separated growth at 48 hours, is shown by the color bar. Here the ratio for ΔLys is shown. The ΔIle ratio plot looks similar with x and y axis transposed. The experimentally mean observed ΔLys ratio for pore sizes of 0.03, 0.1, and 0.2 μm was 0.993 (blue dot next to color bar) and the ΔIle ratio was 1.37 (green dot next to color bar). Thresholds equal to twice the experimental statistic standard deviation (0.314 for ΔLys: blue line next to color bar, and 0.105 for ΔIle: green line) were used to approximate the posterior distribution of the leakages (histograms, black – prior distribution, blue – posterior distribution). **(B)** Example growth curves for different regions of the parameter space.

**Supplemental Figure S7.2:**
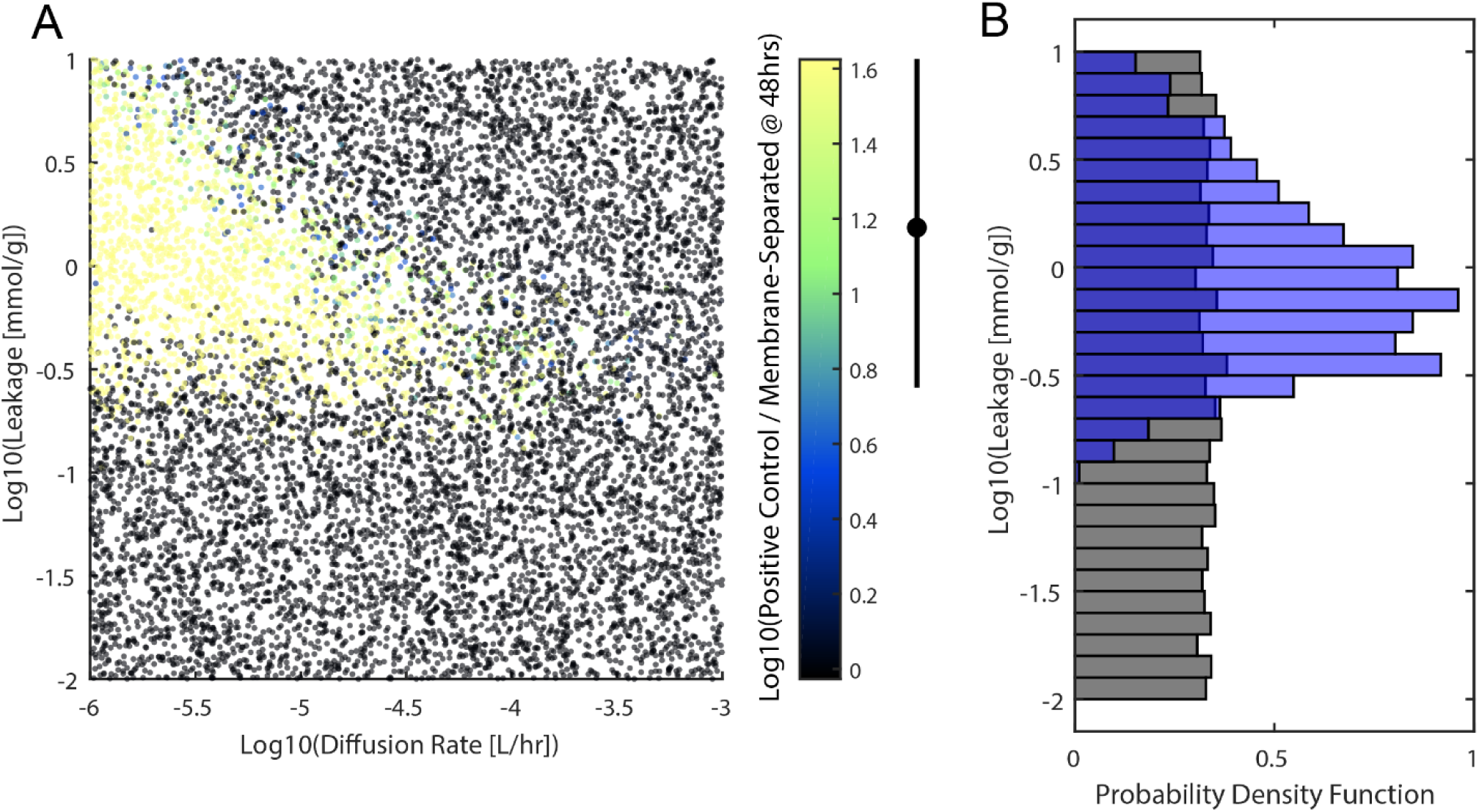
Sampling of diffusion and leakage parameters with noise. All previously fixed parameters (excluding volume) were randomly sampled from a log uniform distribution that varied by 1 order of magnitude around the originally fixed literature estimated value. **(A)** Sampled space of diffusion and leakage parameters. **(B)** Applying an approximate Bayesian computation approach the posterior distribution of the leakage parameter was determined (black – prior distribution, blue – posterior distribution).

**Supplemental Figure S7.3:**
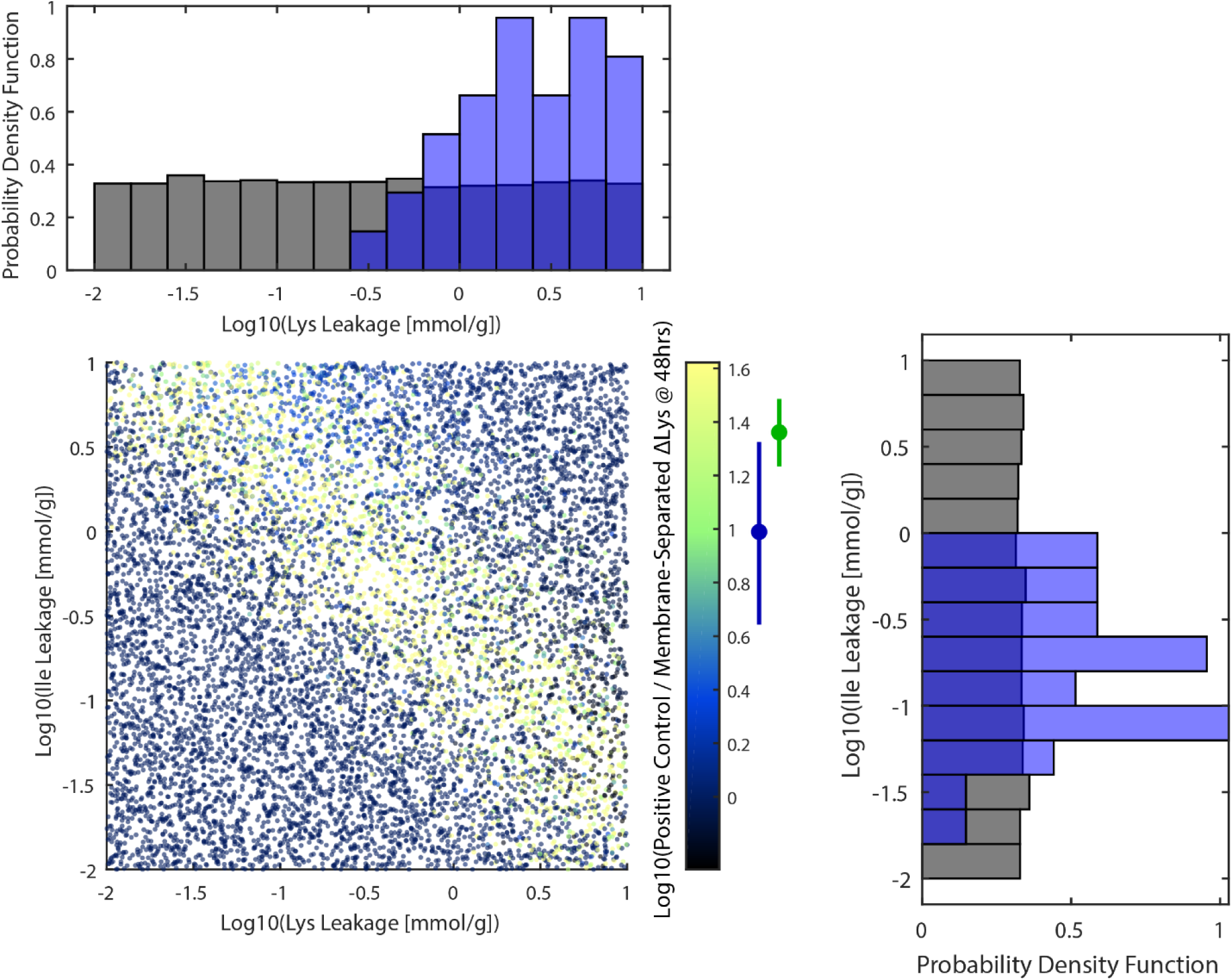
Sampling of unequal leakage parameters with noise. All previously fixed parameters (excluding volume) were randomly sampled from a log uniform distribution that varied by 1 order of magnitude around the originally fixed literature estimated value. The sampled space of unequal leakage parameters is shown. Applying an approximate Bayesian computation approach the posterior distribution of the leakage parameters were determined (black – prior distribution, blue – posterior distribution).

